# Sperm acrosome overgrowth and infertility in mice lacking chromosome 18 pachytene piRNA

**DOI:** 10.1101/2020.09.29.318584

**Authors:** Heejin Choi, Zhengpin Wang, Jurrien Dean

**Author notes:** Correspondence (H.C.), (J.D.).

## Abstract

piRNAs are germline-specific, small non-coding RNAs required to maintain genome integrity and preserve RNA homeostasis during male gametogenesis. In murine adult testes, the highest levels of piRNAs are present in the pachytene stage of meiosis, but their mode of action and function remains incompletely understood. We previously reported that BTBD18 binds to 50 pachytene piRNA-producing loci. Here we show that spermatozoa in gene-edited mice lacking a BTBD18 targeted pachytene piRNA cluster on Chr18 have severe sperm head dysmorphology, poor motility, impaired acrosome exocytosis and are sterile. The absence of Chr18 piRNA results in an imbalance of heat shock proteins associated with renaturing proteins and the ubiquitin-proteasome system involved with protein degradation. The mutant phenotype arises from aberrant formation of proacrosomal vesicles, distortion of the *trans*-Golgi network, and up-regulation of GOLGA2 associated with acrosome dysgenesis. Collectively, our findings reveal central role of pachytene piRNAs in controlling spermiogenesis and male fertility.

## Introduction

Genome integrity and RNA homeostasis are essential for mammalian gametogenesis and rely on piRNAs (P-element induced wimpy testis [PIWI]-interacting RNAs), the most abundant population of small, non-coding RNAs in male gonads. During mouse spermatogenesis, germ cells produce pre-pachytene piRNAs derived from transposable elements and subsequently generate pachytene piRNA from genetically distinct loci scattered throughout the genome^1–3^. The most well-studied and conserved function of pre-pachytene piRNAs is repression of transposons to ensure integrity of the germline genome^4–10^. Pachytene piRNAs derive their designation by expression during meiosis and are considerably more abundant than pre-pachytene piRNAs^11^. Although mice lacking proteins required for pachytene piRNA biogenesis have spermatogenic arrest and male sterility^3, 12–15^, the functions of pachytene piRNAs themselves are much less understood. Hypotheses for their role include: 1) cleaving mRNAs necessary for meiotic progression; and 2) directed degradation of target mRNA analogous to miRNA function in somatic cells^16–19^. We previously identified a pachytene germ cell nuclear protein, BTBD18, that acts as a licensing factor for RNA polymerase II elongation at fifty pachytene piRNA sites scattered across somatic chromosomes. About half of these sites are transcribed on alternative strands of DNA from bi-directional, A-MYB and BTBD18-binding promoters. The absence of BTBD18 in *Btbd18^Null^* mice disrupts piRNA biogenesis, arrests spermiogenesis at an early stage and results in male sterility^20^.

Mouse spermatogenesis has three distinct phases: 1) mitotic proliferation and differentiation; 2) meiosis with two reductive divisions to form haploid gametes; and 3) spermiogenesis in which terminally differentiated, round spermatids undergo a remarkable transformation. During this latter process, male germ cells shed a cytoplasmic droplet and transmogrify into elongated mature spermatozoa with a sperm-unique acrosome overlying a condensed nucleus, a mid-piece filled with mitochondria and a flagellum for forward motility necessary to ascend the female reproductive tract and fertilize eggs. The 16 stages of spermiogenesis can be divided into four phases: Golgi (stages 1-3), cap (stages 4-7), acrosome (stages 8-12), and maturation (stages 13-16)^21–25^. The acrosome is a specialized subcellular, membranous organelle located at the anterior portion of the sperm head. It is an exocytotic vesicle that contains enzymes essential for fertilization, dispersion of cumulus cells and/or sperm penetration of the zona pellucida^21, 22, 26, 27^. Acrosome biogenesis begins in the concave region of the spermatid nucleus during the Golgi phase of spermiogenesis. Golgi-derived proacrosomal vesicles (PVs) accumulate and a single large acrosomal granule (AG) is formed by fusion of small vesicles. The AG attaches to the nuclear envelope via the acroplaxome (Apx), a structure that lies between the inner membrane of the acrosome and nucleus^23^. By combining with additional Golgi-derived vesicles, the size of the acrosome increases and spreads over the anterior nuclear pole during the cap phase of spermiogenesis. The subsequent elongation phase which forms mature spermatozoa is mediated by the perinuclear ring of the manchette and its associated microtubules that are subsequently degraded^24^. Despite the well-documented morphologic changes in acrosome biogenesis during spermiogenesis, the underlying molecular mechanisms remain to be determined.

Because of chromatin condensation in which nuclear histones are replaced with disulfide-bond rich protamines, elongating spermatids and mature sperm are transcriptionally silent. Thus, accurate post-transcriptional quality control of RNA and proteins is critical for normal spermatogenesis. To explore the functional importance of pachytene piRNAs, we have used CRISPR/Cas9 genome editing to establish mouse lines unable to express specific piRNA precursors from a highly prolific^28^, bi-directional pachytene piRNA clusters on Chromosome 18 (Chr18). By investigating proteome and transcriptome profiles of these non-proliferating cells, we have discovered that Chr18 pachytene piRNAs maintain protein homeostasis and ensure proper acrosome formation. Up-regulated transcripts and proteins have been identified including *Golga2* associated with acrosome overgrowth. Taken together our data indicate that Chr18 pachytene piRNA plays an essential role during spermiogenesis, which is critical for male fertility.

## Results

### Chr18 pachytene piRNAs are required for spermatogenesis and male fertility

To disrupt the bi-directional piRNA promoter at the precursor locus on Chr18, we used a pair of single-guide RNAs (Fig. 1a, Extended Data Fig. 1a, c). From founders that passed the mutant allele through their germline, we established mouse lines with a ∼1.3 kb deletion in the promoter of the Chr18 piRNA clusters and bred them to homozygosity (referred to as *Chr18^Δ/Δ^*, Extended Data Fig. 1b, c). To determine if the loss of pachytene piRNAs transcribed on Chr18 affected reproduction, we mated C57BL/6 females with *Chr18^Δ/Δ^* or control (*Chr18^+/Δ^*) male mice. Vaginal plugs were observed in all females. *Chr18^Δ/Δ^* male mice did not produce litters whereas control males did and *Chr18^Δ/Δ^* female mice had normal fertility (Fig. 1b).

**Fig. 1.**
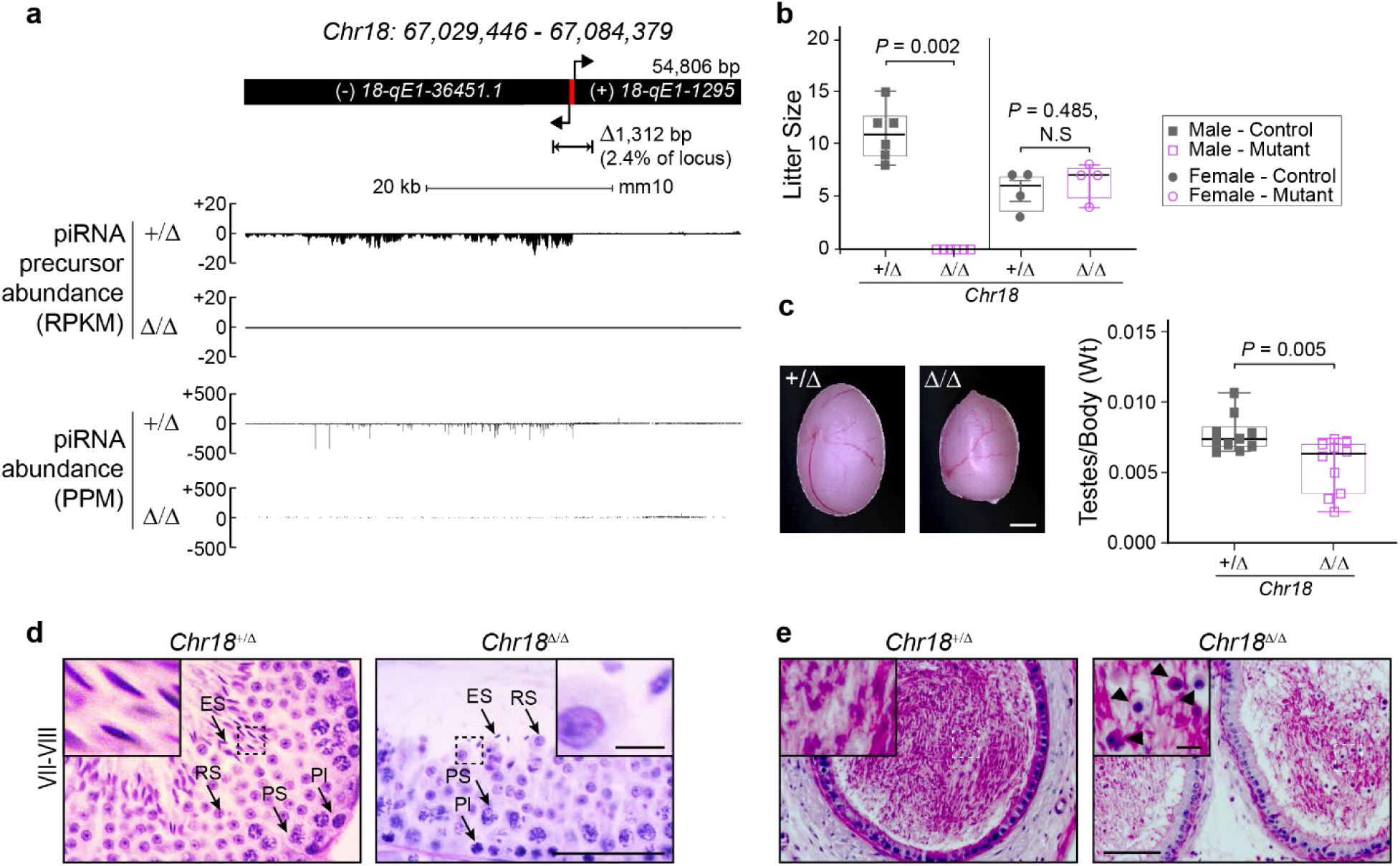
Pachytene piRNA derived from Chr18 is essential for spermatogenesis and male fertility. **a**, Schematic diagram of the Chr18 piRNA coding region and the DNA fragment deleted using CRISPR/Cas9 to generate bi-directional promoter deletion (Δ/Δ) mutant mice (top). See also Extended Data Fig. 1a-c and Extended Table 3, 4. Red bar, A-MYB and BTBD18 binding loci. Precursor and mature piRNA abundance at Chr18 piRNA cluster in P28 testes of heterozygous (+/Δ) and homozygous (Δ/Δ) mutant mice (bottom). RPKM, reads per kilobase million; PPM, parts per million reads. **b**, Average litter size of adult male (squares, *n* = 6) and female (circles, *n* = 4) control and mutant mice. **c**, Representative macroscopic appearance (left) and quantification (right) of testes weight of 8 wk/old control and mutant mice (*n* = 10 per each genotype). Scale bar, 2 mm. **d**, Testicular sections from 8 wk/old mice were stained with periodic acid-Schiff (PAS) and hematoxylin (H) (*n* = 3). Stage of seminiferous epithelium cycles was determined by morphology of spermatocytes and rounds spermatids. Pl, preleptotene spermatocyte; PS, pachytene spermatocyte; RS, round spermatid; ES, elongating spermatid. Scale bar, 50 μm; inset, scale bar, 5 μm. **e**, PAS&H staining of cauda epididymis from 12 wk/old mice (*n* = 2). Black arrow heads, sloughing germ cells. Scale bar, 50 μm; inset, scale bar, 5 μm. **b**, **c** The box indicates median ± interquartile range, the whiskers indicate the highest/lowest values and midlines are median values. Two-tailed *P* values were calculated using Student’s *t*-test.

The growth rates of *Chr18^Δ/Δ^* and control male mice did not differ, but the average weight of testes from adult mutant mice was ∼45% less than controls (Fig. 1c). Because *Chr18^Δ/Δ^* male mice were sterile, we investigated the stage at which spermatogenesis failed. Compared to *Chr18^+/Δ^* controls, spermatocytes, round spermatids, and early elongating spermatids were largely unaltered, while condensed spermatids at steps 14-16 of spermiogenesis (stages III-VIII of spermatogenesis) were significantly reduced in the seminiferous tubules of *Chr18^Δ/Δ^* testes (Fig. 1d). Concomitantly, there was a significant increase in the number of apoptotic cells in seminiferous tubules of *Chr18^Δ/Δ^* mice (Extended Data Fig. 2a). In addition, we frequently found histological abnormalities including disordered arrangement of elongating spermatids and vacuolation in *Chr18^Δ/Δ^* testes (Extended Data Fig. 2b, c). *Chr18^Δ/Δ^* mice were infertile, but elongating spermatozoa associated with sloughed germ cells were present in the lumen of their epididymides (Fig. 1e, Extended Data Fig. 2d). To further characterize the pathology, sperm were collected from the vas deferens and cauda epididymis of controls and *Chr18^Δ/Δ^* mice. The total number of sperm from *Chr18^Δ/Δ^* mice was significantly less than obtained from controls (Fig. 2a). Based on computer-assisted sperm analysis (CASA), caudal sperm from *Chr18^Δ/Δ^* mice had decrease progressive motility and all parameters describing the speed of their movements, including path velocity (VAP), track velocity (VCL), and linear velocity (VSL) were significantly reduced (Fig. 2b-d). In addition, most sperm from *Chr18^Δ/Δ^* mice exhibited various abnormalities of acrosomal overgrowth (Fig. 2e).

**Fig. 2.**
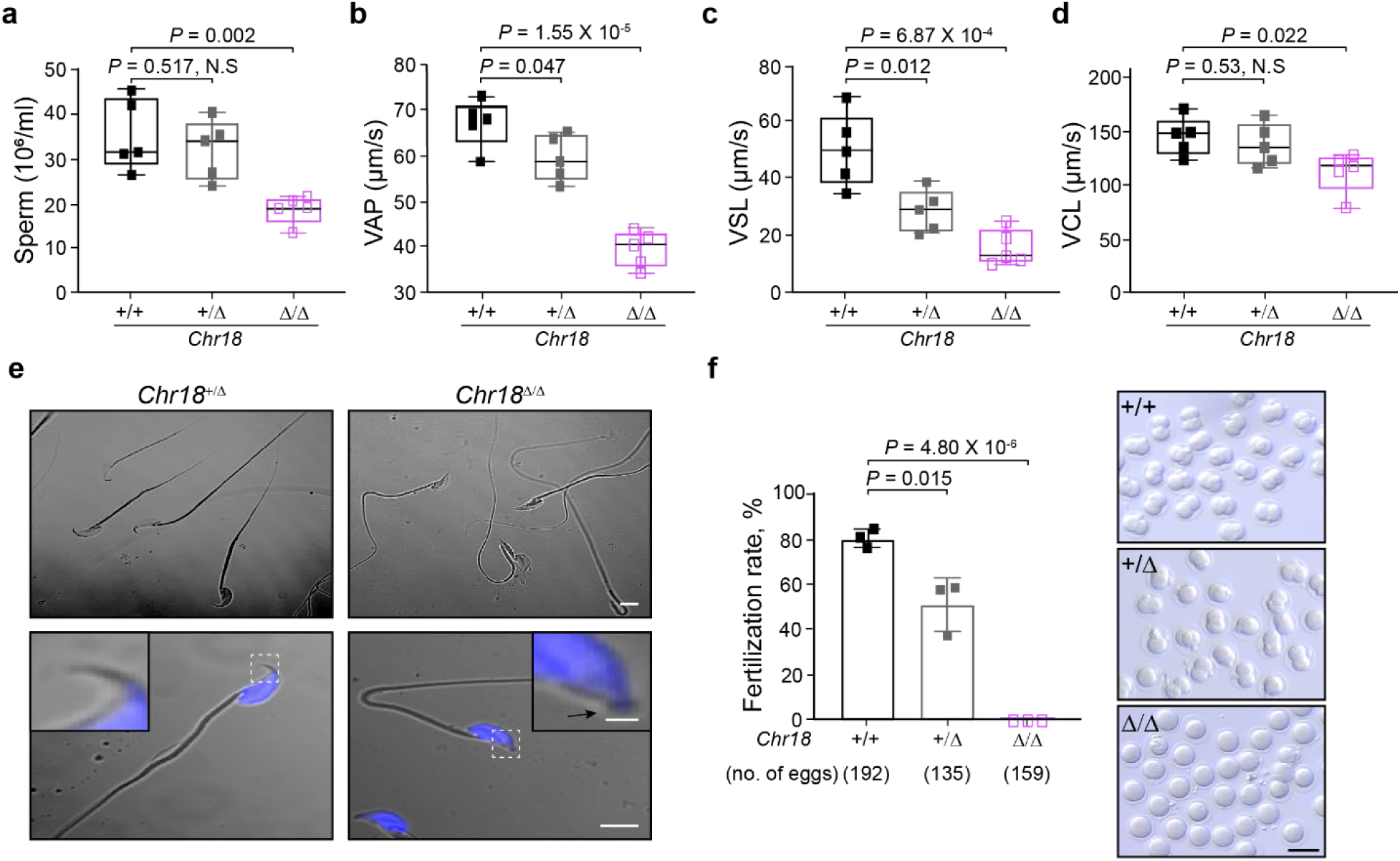
Impaired motility and *in vitro* fertilization with *Chr18^Δ/Δ^* sperm. **a**, Box plots showing sperm counts (*n* = 5) and **b-d**, Computer-Assisted Sperm Analysis (CASA) assay of sperm motility from 8 wk/old mice (*n* = 5 for each genotype). Path velocity (velocity average path, VAP) (**b**), track velocity (velocity curvilinear, VCL) (**c**), and linear velocity (velocity straight line, VSL) (**d**) of spermatozoa from *Chr18^+/+^*(black square), *Chr18^+/Δ^* (grey square), and *Chr18^Δ/Δ^* (purple square) mice. **e**, Representative differential interference contrast (DIC) micrograph images of sperm from 12 wk/old mice, with nuclei counterstained with Hoechst 33342 (blue) (*n* = 3). Scale bar, 5 μm; inset, scale bar, 0.5 μm; Arrow, apical hook. **f**, *Chr18^Δ/Δ^* sperm are unable to fertilize wildtype eggs *in vitro* (left). The average fertilization rate (mean ± s.d.) from three independent experiments is presented. Each square represents individual male mice that were used for IVF. The total number of analyzed eggs per condition is in the parenthesis. The appearance of two-cell-stage embryos (right) determined the fertilization rate. Scale bar, 100 μm. **a-d**, **f** The box indicates median ± interquartile range, the whiskers indicate the highest/lowest values and midlines are the median values. Two-tailed *P* values were calculated using Student’s *t*-test.

*In vitro* fertilization (IVF) assays were performed to investigate the ability of *Chr18^Δ/Δ^* sperm to fertilize eggs. Mature eggs obtained from wild-type females were inseminated with capacitated sperm from *Chr18^Δ/Δ^* and controls. After 24 hours, fertilization rates were determined by the presence of two-cell embryos. Whereas 156 of 192 (81.3%) and 68 of 135 (50.4%) eggs were fertilized by sperm derived from *Chr18^+/+^*and *Chr18^+/Δ^* sperm, respectively, no fertilization was observed with *Chr18^Δ/Δ^* sperm (Fig. 2f). Notably, although IVF was performed in media containing reduced glutathione to destabilize the extracellular zona pellucida (ZP)^29, 30^, eggs were not fertilized by *Chr18^Δ/Δ^* sperm. The IVF results documented that *Chr18^Δ/Δ^* sperm can pass through the cumulus cell layers but fail to penetrate the zona matrix. Taken together, the defects *in vivo* and *in vitro* of *Chr18^Δ/Δ^* sperm support an essential role of Chr18 pachytene piRNAs in promoting male fertility.

### Defects of spermiogenesis and acrosome exocytosis in *Chr18^Δ/Δ^* mice

Scanning (SEM) and transmission electron microscopy (TEM) of sperm heads, acrosomes and cross-sections of tails were used to more precisely define sperm dysmorphology. The majority of *Chr18^Δ/Δ^* sperm heads were abnormally round shaped with shortened apical hooks, smaller apical angles and bulges in the acrosome region (Fig. 3a, b). Although all layers, including plasma, outer and inner acrosomal as well as nuclear membranes were intact, the dramatic overgrowth of the acrosome excessively folded onto itself (Fig. 3c). Cross sections of the mid-piece of *Chr18^Δ/Δ^* sperm document a well-defined mitochondrial sheath, normally arranged outer dense fibers (ODF) and an axoneme with an intact “9+2” microtubule structure. However, the axonemal complex was abnormal, and the outer dense fibers were unassembled in the principal piece (tail) of *Chr18^Δ/Δ^* sperm which could account for the lost motility (Fig. 3d).

**Fig. 3.**
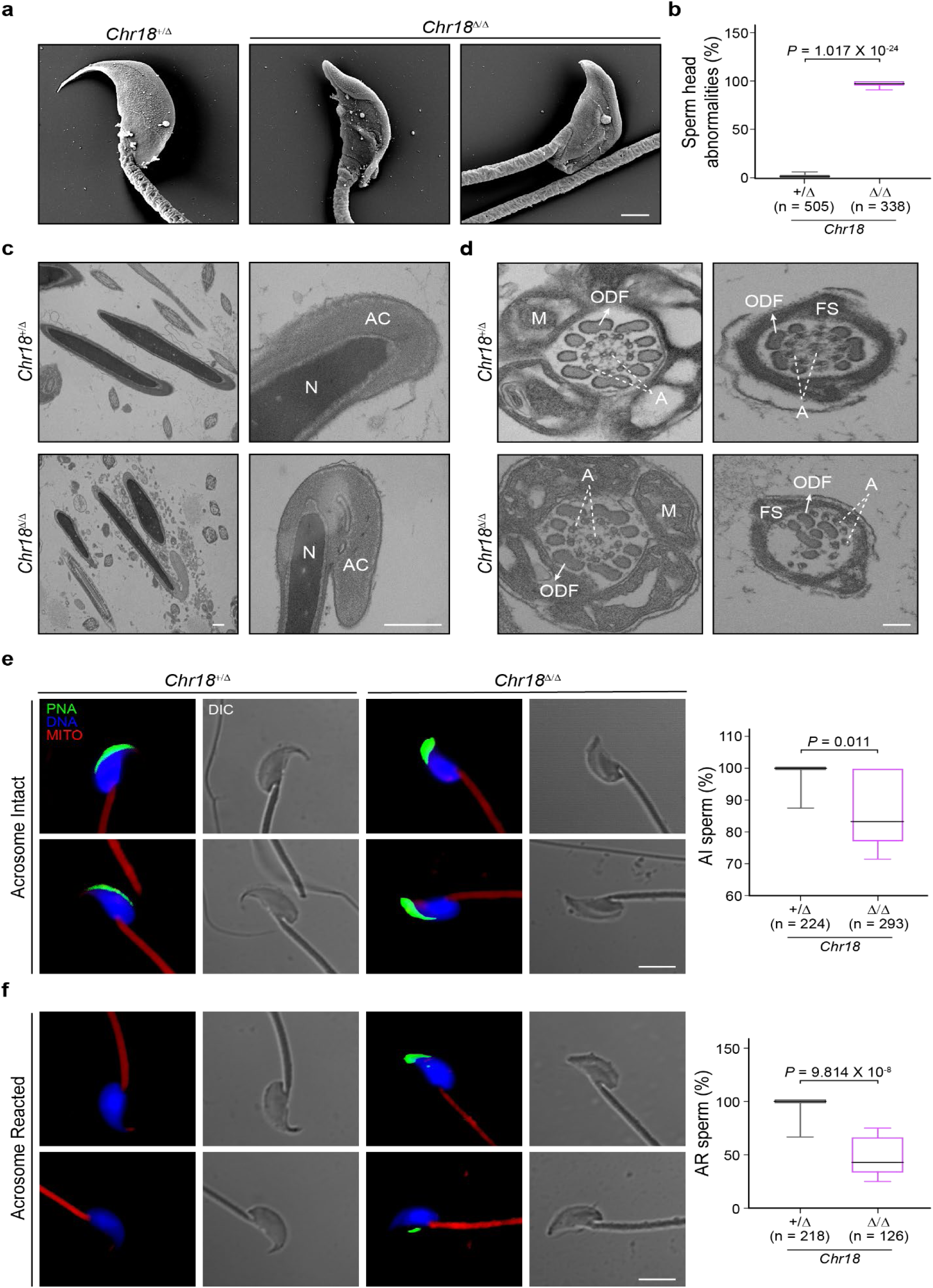
Defective spermiogenesis and impaired acrosome reaction in *Chr18^Δ/Δ^* mice. **a**, **b**, Representative scanning electron microscopy (SEM) images (**a**) and quantification (**b**) of malformed sperm head in 12 wk/old control and *Chr18^Δ/Δ^* mutant mice (*n* = 3 per group). Scale bar, 1 μm. **c**, Transmission electron micrographs (TEM) of sperm heads from 12 wk/old mice (*n* = 4 for each genotype). Scale bar, 0.5 μm; AC, acrosome; N, nucleus. **d**, TEM images of sperm mid (left) and principal (right) pieces. Scale bar, 0.5 μm; ODF, outer dense fiber; M, mitochondria; A, 9+2 axoneme; FS, fibrous sheath. **e**, **f**, Representative confocal images (left) and quantification (right) of acrosome intact (**e**) and acrosome reacted (**f**) cauda epididymal sperm from 12 wk/old control and mutant mice. Acrosome exocytosis induced with calcium ionophore A23187. Sperm were stained for fluorescent-dye-labeled peanut agglutinin (PNA) (acrosome, green); MitoTracker Red FM (mitochondria, red); Hoechst 33342 (DNA, blue). Scale bar, 5 μm; AI, acrosome intact; AR, acrosome reacted; DIC, differential interference contrast (*n* = number of sperm total from 3 independent experiments). **b**, **e-f** The box indicates median ± interquartile range, the whiskers indicate the highest/lowest values and midlines are median values. Two-tailed *P* values were calculated using Student’s *t*-test.

To further investigate the inability of *Chr18^Δ/Δ^* sperm to fertilize eggs, we used Alexa Fluor 488-conjugated peanut agglutin (PNA) to determine acrosome exocytosis which is a prerequisite for gamete fusion^21^. PNA binds to the outer acrosomal membrane and, in agreement with previous results^31–33^, fluorescent staining on the crescent region of acrosome-intact *Chr18^+/Δ^* control sperm disappeared after induction of the acrosome reaction by calcium ionophore, A23187. The dorsal edge of the acrosome is not fully elongated in *Chr18^Δ/Δ^* sperm and, consistent with SEM and TEM images, exhibited accumulated fluorescence on the acrosomal vesicle bulge. Notably, about half of *Chr18^Δ/Δ^* sperm had slightly reduced PNA signal on their acrosomes which did not disappear after induction of acrosome exocytosis (Fig. 3e, f). Taken together, the differences in sperm velocities and impaired acrosomal reaction contribute to the inability of *Chr18^Δ/Δ^* sperm to fertilize wildtype eggs.

### Altered acrosome formation in *Chr18^Δ/Δ^* spermatids

To link development of acrosome abnormalities to the phase of spermiogenesis, we used light microscopy to examine spermatogenic cells from control and *Chr18^Δ/Δ^* mice. The Golgi apparatus of spermatids consists of several stacks of saccules with a *cis*-network facing the endoplasmic reticulum (ER), and a *trans*-Golgi network (TGN) facing the nuclear envelope. Budding proacrosomal vesicles (PVs) from the TGN initiate acrosome formation and proper trafficking from the TGN toward the nucleus is essential for normal shaping and sizing of the acrosome. Periodic acid-Schiff (PAS) and PNA staining documented the presence of normal-shaped acrosomes in the Golgi phase of *Chr18^Δ/Δ^* spermatids and there were no obvious differences with control spermatids (Extended Data Fig. 3a, b, upper panels).

However, using TEM in control mice, we observed umbrella shaped TGNs and multiple PVs of uniform size located between the TGN and the nuclear membrane in control spermatids. Although PVs were present in *Chr18^Δ/Δ^* spermatids, they were not uniform in size and appeared larger than those in control sperm. Moreover, we frequently observed that the lamellar structure of the TGN formed loose whorls in *Chr18^Δ/Δ^* spermatids (Fig. 4a). These observations suggest that there are abnormalities in the vesicles budding from the TGN in *Chr18^Δ/Δ^* spermatids. To quantify the foregoing observations, we counted and measured the diameter of PVs on the TEM sections (Extended Data Fig. 4a). C*hr18^Δ/Δ^* spermatids tend to produce more PVs with larger diameters than vesicles in control spermatids (Fig. 4b, c). These results indicate that aberrant PV formation and budding from the TGN resulted in formation of deformed acrosomes in *Chr18^Δ/Δ^* spermatids.

**Fig. 4.**
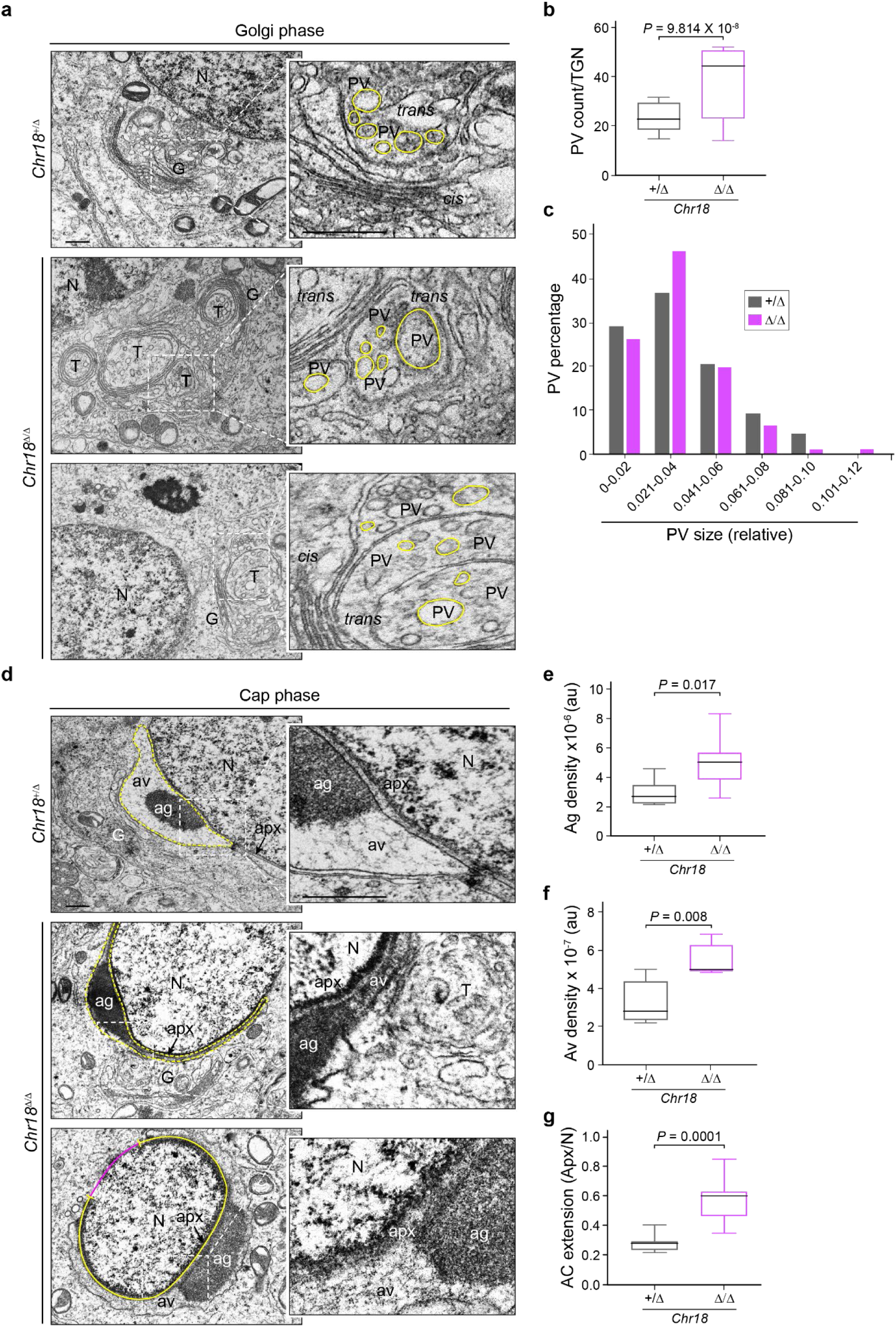
Deformed acrosome formation from disrupted proacrosomal vesicles in *Chr18^Δ/Δ^* mice. **a**, **b**, Representative TEM images (**a**) and quantification (**b**) of proacrosomal vesicles in Golgi phase round spermatids in 12 wk/old control and mutant mice (*n* = 2 per group). Enlarged insets (right) correspond to the dashed box (left) of the Golgi apparatus (**a**). Scale bar, 0.5 μm; G, Golgi apparatus; T, *trans*, and TGN, *trans*-Golgi network; *cis*, *cis*-Golgi network; PV, proacrosomal vesicle; N, nucleus. **c**, PV size distribution determined by measuring diameters in control and mutant round spermatids at the Golgi phase (**a**). The diameters are presented in relative units. PV sizes are binned as indicated (see Extended Data Fig. 4a). **d-g**, TEM images (**d**) of cap phase round spermatids in 12 wk/old control and mutant mice (*n* = 2 for each genotype). The acrosome at cap phase round spermatids in dashed boxes (left) are enlarged (right). Scale bar, 0.5 μm; Ac, acrosome; Ag, acrosomal granule; Av, acrosomal vesicle; Apx, acroplaxome. Quantification of acrosomal granule density (**e**), acrosomal vesicle density (**f**) and acrosome enlargement (**g**). AU, arbitrary unit. See Extended Data Fig. 4b. **b**, **e-g** The box indicates median ± interquartile range, the whiskers indicate the highest/lowest values and midlines are median values. Two-tailed *P* values were calculated using Student’s *t*-test.

As spermiogenesis proceeded to the cap phase, the defects in *Chr18^Δ/Δ^* spermatids became more obvious. In the cap phase, proacrosomal vesicles (PV) fuse with each other to form the acrosome granule (AG) at the acroplaxome (Apx) which anchors the acrosome (Ac) to the nuclear membrane over which the acrosome flattens^34^. In control spermatids stained with PNA, the acrosome grew into a single cap-like structure that covered nuclei (Extended Data Fig. 3a, 3b, middle panels). In contrast, *Chr18^Δ/Δ^* spermatids displayed a notably enlarged acrosomal cap composed of highly electron dense acrosomal granules and vesicles. Aberrant PV formation in the Golgi phase inundated the acrosome with vesicles and, as determined by the marginal ring of the Apx, the nucleus became covered with an overgrown acrosome at the cap phase in *Chr18^Δ/Δ^* spermatids (Fig. 4d-g, Extended Data Fig. 4b). In the subsequent acrosome phase of spermiogenesis, plump and thickening acrosomes were observed in *Chr18^Δ/Δ^* spermatids but positioning of the perinuclear ring of the manchette^24^ was comparable to controls (Extended Data Fig. 3a, 3b, lower panels; Fig. 4c). Together, these results indicate that Chr18 pachytene piRNA plays an essential role in regulating acrosome biogenesis during spermiogenesis.

### Disruption of protein homeostasis in *Chr18^Δ/Δ^* mice

According to the foregoing observations, loss of Chr18 pachytene piRNA could result in defective protein homeostasis during spermiogenesis. To investigate the molecular basis of the sperm dysmorphology, we analyzed proteomic profiles of testicular cells and mature sperm from control and *Chr18^Δ/Δ^* mice (Extended Data Table 1, 2). Compared with *Chr18^+/Δ^* controls, we identified 168 and 291 differentially expressed proteins (DEP) from testes and mature sperm, respectively, in *Chr18^Δ/Δ^* mice (*P* < 0.05, Fig. 5a, Extended Data Fig. 5a). Ingenuity Pathway Analysis (IPA) of the 62 (testes) and 220 (sperm) DEPs more abundant in *Chr18^Δ/Δ^* than control mice, predicted defects in protein homeostasis, including post-translational modifications, protein folding and aggregation. Interestingly, IPA of these proteins also projected down-regulation in the quantity of germ cells, sperm disorders, azoo- or oligozoospermia, decreased testicular mass, degeneration and defects in reproductive system development compared with *Chr18^+/Δ^* controls (Fig. 5b). Consistent with these predictions, mass spectrometry documented significant increases in ODF2 (Outer dense fiber of sperm tails 2), CHIP/STUB1 (C-terminus of HSC70-interacting protein), BAG5 (Bcl-2-associated athanogene 5), and BOLL (Boule homolog) in *Chr18^Δ/Δ^* mice (Fig. 5a, c). ODF2 is a major component of outer dense fibers (ODFs). The ODFs are prominent sperm tail-specific cytoskeletal structures and are thought to be contractile in the sperm tail^35, 36^. Indeed, the absence of ODF2 resulted in abnormal motility and bent tails, but the ultrastructure of *Odf2^Null^* spermatozoa displayed marginal defects in mid- and principal pieces^37^. In contrast, increased expression of ODF2 (Fig. 5d) results in poor motility and severely deformed structural defects in *Chr18^Δ/Δ^* sperm (Fig. 2b-d, 3d). STUB1 is a ubiquitously expressed cytosolic E3-ubiquitin ligase that regulates HSP70-mediated protein degradation^38^. BAG5, a member of a larger family of proteins, directly interacts with HSP70 and inhibits HSP70-mediated refolding of misfolded proteins^39^. Both STUB1and BAG5 are related to pathways of cellular response to stress and protein kinase and chaperone binding to target proteins for degradation^38–41^. In addition, it has been reported that diseases associated with BOLL include azoospermia and male infertility^42–44^.

**Fig. 5.**
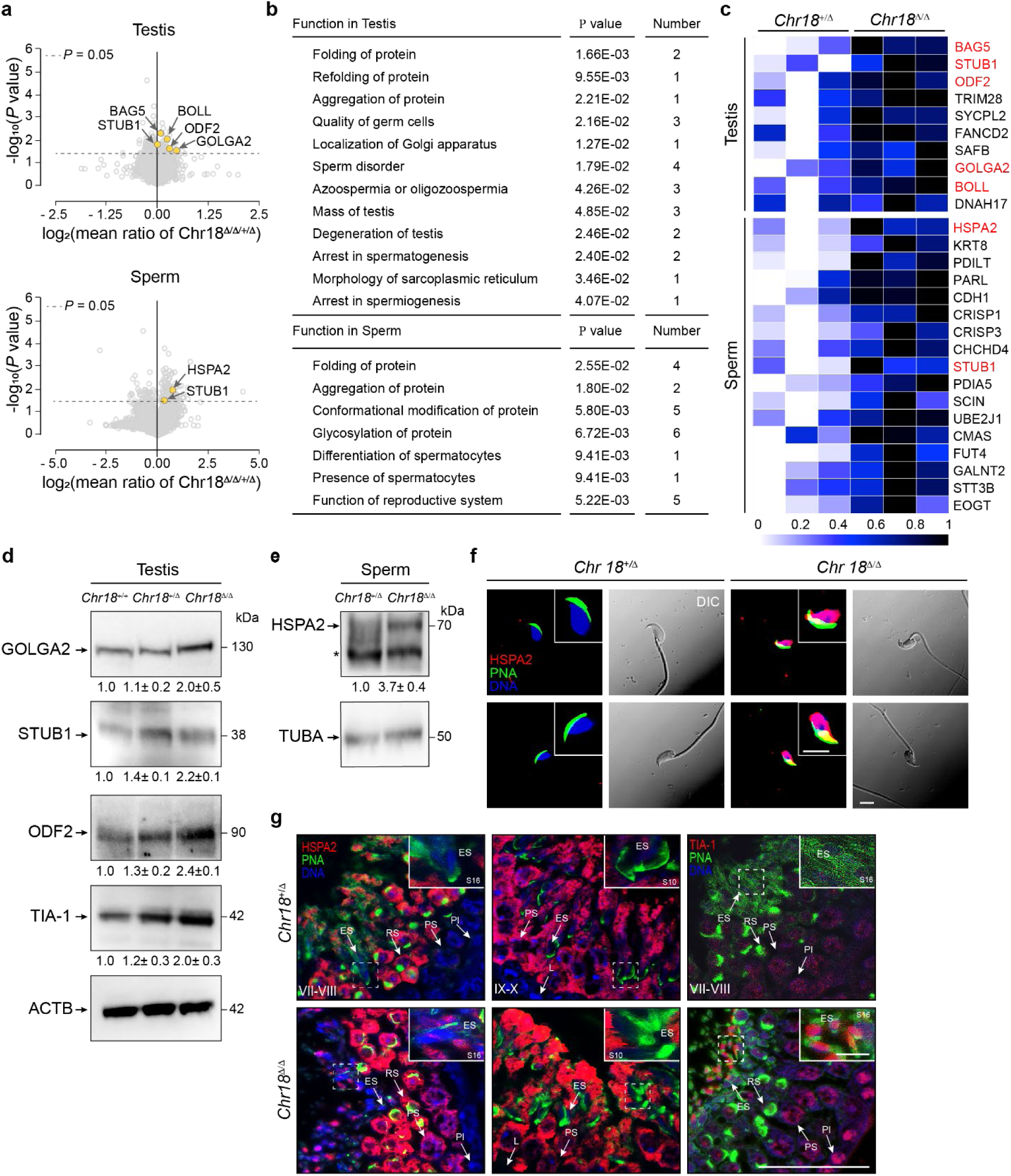
An increased abundance of proteins in *Chr18^Δ/Δ^* mice. **a**, **b**, TMT-labeling mass spectrometry volcano plots (**a**) showing enrichment values and corresponding significance levels for proteins in testis (top) and sperm (bottom) in 12 wk/old control and mutant mice (*n* = 3 for each genotype). Colored dots are significant proteins in the indicated pathways in table (**b**). The dashed horizontal line indicates two-tailed *P* = 0.05. **c**, Heat map of representative proteins from pathways listed in (**b**) in testis (top) and sperm (bottom). Significant proteins in pathways **(b)** are highlighted in red. See also Extended Table 1, 2. **d**, **e**, Immunoblot of GOLGA2, STUB1, and ODF2 in testes from *Chr18^+/+^*, *Chr18^+/Δ^*, and *Chr18^Δ/Δ^* (**d**) and HSPA2 in sperm from *Chr18^+/Δ^* and *Chr18^Δ/Δ^* (*n* = 3 for each genotype) (**e**). ACTB (**d**) and TUBA (**e**) were used as loading controls. Quantification of band intensity for indicated proteins is shown as the relative expression compared with *Chr18^+/+^*(**d**) and *Chr18^+/Δ^* (**e**) controls. *, non-specific. Data were generated from three independent experiments. **f**, Representative confocal images of cauda epididymal sperm from 12 wk/old control (left) and mutant mice (right). On the right of the paired images, sperm were stained for HSPA2 (red); fluorescent-dye-labeled peanut agglutinin (PNA) (acrosome, green); Hoechst 33342 (DNA, blue). The left of the paired images was obtained with differential interference contrast (*n* = 3). Scale bar, 5 μm. **g**, Representative confocal microscopic images of testis sections from 8 wk/old control and mutant mice (n=3) stained for HSPA2 (red, right, middle), TIA-1 (red, left), PNA (acrosome, green) and Hoechst 33342 (DNA, blue). Pl, preleptotene; L, leptotene; PS, pachytene spermatocyte; RS, round spermatid; ES, elongating spermatid. Scale bar, 50 μm; inset, scale bar, 5 μm. Stages of seminiferous epithelium cycles were determined by morphology of spermatocytes and round spermatids with PNA staining.

We observed significantly increased abundance of STUB1 in *Chr18^Δ/Δ^* testes compared to controls (Fig. 5d). Moreover, mass spectrometry detected increased abundance of STUB1 and heat shock protein 2 (HSPA2) in *Chr18^Δ/Δ^* mature sperm. Decreased expression of HSPA2 in meiotic and post-meiotic germ cells has been reported in normal mice^45–47^, but we observed HSPA2 retention in *Chr18^Δ/Δ^* mature sperm (Fig. 5e, f). To substantiate this observation, we determined by immunofluorescence that the HSPA2 nuclear signal persisted in late stage spermatids and was detected in a stage-specific manner at step 16 of spermiogenesis (stages VII-VIII of spermatogenesis) of *Chr18^Δ/Δ^* mice (Fig. 5g, Extended Data Fig. 5d). HSPA2 is a testis-specific member of the heat shock protein 70 (HSP70) family. Heat shock proteins (HSPs) act as molecular chaperones that protect proteins from aggregation, catalyze the proper folding of nascent and mis-folded proteins and facilitate degradation under physiological or stressful conditions^48^. The ubiquitin-proteasome system (UPS) is an error-checking system that is invoked for destruction of improperly or unnecessarily synthesized proteins^49^. HSPs and UPS systems have evolved to assist renaturing (HSPs) or destroying (UPS) proteins damaged from stress and collaboration between these systems is essential to triage protein quality (Extended Data Fig. 6a)^50^. Thus, proteins essential for protein triage have undergone significant changes in abundance in *Chr18^Δ/Δ^* mice which implies critical roles for Chr18 piRNAs in protein homeostasis beyond acrosome biogenesis.

Although STUB1 and BAG5 inhibit the chaperon function of HSP70, the amounts of chaperone and proteases increased in parallel which suggests a competitive balance between them in *Chr18^Δ/Δ^* mice. Furthermore, protein overproduction and acrosomal overgrowth reflect perturbed balance of protein triage and, as previously suggested^50^, can result in increased sensitivity to stress in *Chr18^Δ/Δ^* mice. BOLL is a component of stress granules (SG) in heat stressed mouse male germ cells^43^. We, therefore, examined the sensitivity of *Chr18^Δ/Δ^* spermatids to stress with cytotoxic granule associated RNA binding protein, TIA-1, a marker for SGs. Consistent with previous reports^43, 51^, TIA-1 was in the nuclei of spermatocytes and round spermatids in control mice. In contrast, while we detected TIA-1 in spermatocytes and round spermatids, we observed unique expression pattern of TIA-1 in step 16 spermatids nucleus and increased level of TIA-1 in *Chr18^Δ/Δ^* testes (Fig. 5d, g). This finding differs from BOLL and TIA-1 relocation patterns to the cytoplasmic SGs of spermatocytes induced by heat stress and, instead resembles HSPA2 expression pattern in *Chr18^Δ/Δ^* mice. Furthermore, HSPA2 and TIA-1 localization in the nucleus, but not all their overall abundance, depends on the stage of spermiogenesis. Taken together, our data suggest that the absence of Chr18 pachytene piRNA resulted in failure of protein quality control and leads to protein aggregation and stress granule formation in the late stages of spermiogenesis.

### Chr18 pachytene piRNAs control *Golga2* mRNA abundance and cleaves in the 3’-UTR

To further investigate molecular consequences in *Chr18^Δ/Δ^* mice, we isolated mRNA and small RNA from controls and *Chr18^Δ/Δ^* testes at P28 and performed RNA-seq. The mRNA abundance in *Chr18^Δ/Δ^* testes was extensively altered compared to *Chr18^+/+^*controls. In contrast, C*hr18^Δ/Δ^* mutants had modest but significant changes in mRNA abundance compared with *Chr18^+/Δ^* controls (Extended Data Fig. 7a-b). RNA-seq analyses identified 8 up-regulated and 5 down-regulated genes in *Chr18^Δ/Δ^* testes compared with *Chr18^+/+^*and *Chr18^+/Δ^* controls (*P* < 0.01, FDR < 0.1, Extended Data Table 3, 4). However, the steady-state abundance of piRNA precursors was unaffected outside the Chr18 piRNA cluster (Extended Data Fig. 7d). We also investigated whether pachytene piRNA elimination forced transposon de-repression in *Chr18^Δ/Δ^* testes. The ordinary abundance of transposon RNA in *Chr18^Δ/Δ^* testes documented that the fertilization and spermiogenic defects do not reflect a failure to silence transposons (Extended Data Fig. 7e, Extended Data Table 6). Together with small RNA-seq (Extended Data Fig. 7f), these analyses document that deletion of the bi-directional promoter at the Chr18 pachytene piRNA cluster eliminates precursor and processed piRNAs encoded at the site without an accompanying effect on other piRNA clusters or transposons.

Despite the severely deformed sperm head morphology, the most enriched transcripts in gene ontology (GO) analysis of *Chr18^Δ/Δ^* mice related to chromosome segregation, DNA repair, and meiotic recombination (Extended Data Fig. 7g, Extended Data Table 5). However, immunostaining of meiotic chromosome spreads of *Chr18^Δ/Δ^* spermatocytes confirmed successful synapsis and recombination in the absence of Chr18 piRNA (Extended Data Fig. 7h). Therefore, we further searched annotated gene function that might be related to the observed head dysmorphology in *Chr18^Δ/Δ^* mice. Among transcripts that were up-regulated in mutant mice, we identified *Golga2* that encodes a *cis*-side localized Golgi matrix protein (Extended Data Fig. 7c). It has previously been reported that the absence of GOLGA2 in gene-edited mice results in male infertility^44^. These mutant mice, lack acrosomes, have round sperm heads and mitochondrial defects similar to human globozoospermia which is opposite to the acrosomal overgrowth phenotype observed in *Chr18^Δ/Δ^* mice. In addition, overexpression of GOLGA2 in heterologous cells results in irregular and incorrectly aligned stacks of abnormally elongated Golgi including bending and horseshoe-like structures which are similar with *Chr18^Δ/Δ^* spermatids^52^. Intriguingly, four piRNAs annotated at the Chr18 site have a seed sequence from the second to fifteenth nucleotides complementarity to the 3’-UTR of *Gogla2* (Fig. 6b) and it has been reported that piRNA-guided MIWI targets *Golga2* by cleaving the 3’-UTR^18^. In re-analyzing our TMT mass spectrometry data, we found a correlation between mRNA and protein expression level of GOLGA2 in *Chr18^Δ/Δ^* testes (Extended Data Fig. 5b) and validated this finding by immunoblot (Fig. 5d). Compared with *Chr18^+/+^*and *Chr18^+/Δ^* controls, the level of GOLGA2 protein was higher in *Chr18^Δ/Δ^* testes and this increase does not reflect a failure of cytoplasmic depletion in *Chr18^Δ/Δ^* spermatids during spermiogenesis (Extended Data Fig. 5c). Previous investigations reported that the absence of GOLGA2 did not affect PV secretion and thus, the piRNAs absent in *Chr18^Δ/Δ^* spermatids must reflect additional molecular defects to account for the observed aberrant PV and TGN.

**Fig. 6.**
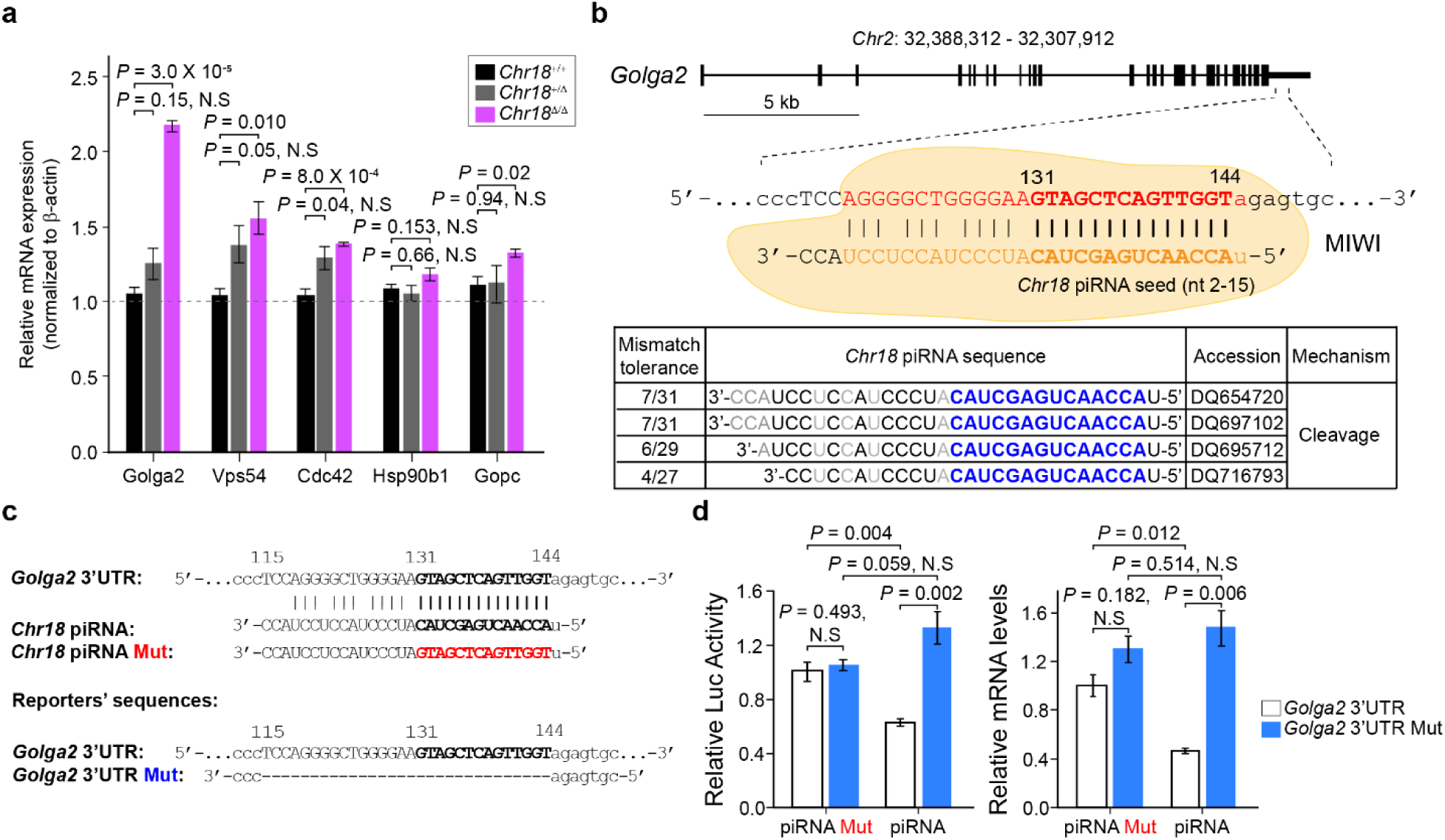
MIWI/*Chr18* pachytene piRNAs cleavage target in the *Golga2* 3’-UTR. **a**, Quantitative RT-PCR validation of up-regulated genes related to acrosome biogenesis using *β-actin* transcript as an internal control (*n* = 3 per genotype). **b**, A schematic showing the targeting of mouse *Golga2* cleavage sites by *Chr18* piRNAs (top) and the best *Chr18* piRNA match to the *Golga2* cleavage site (middle). The blue and grey sequence denotes complementary and mismatched sequence, respectively. Each vertical bar denotes a perfect base pair between the cleavage site and the guide piRNA that can best complement each cleavage site is shown in the table (bottom). **c**, Predicted *Chr18* piRNA regulatory elements in the *Golga*2 3’-UTR, and the sequences of synthetic piRNA, piRNA Mut (negative control) (top), and deletion mutated 3’-UTR-*Renilla* luciferase reporters (bottom). **d**, Dual luciferase reporter assays of the effect of Chr18 piRNA on reporter activities with cognate piRNA Mut, as a negative control (left). Quantitative RT-PCR analyses of the effect of Chr18 piRNA on reporter mRNA levels (right). **a**, **d** Data are shown as mean ± s.d. Two-tailed *P* values were calculated using Student’s *t*-test. Results shown reflect three independent experiments.

These data suggest that piRNAs from Chr18 target genes to ensure control of mRNA abundance during acrosome biogenesis that occurs in mouse spermiogenesis. Both transcriptome and proteome analyses documented that loss of Chr18 piRNA does not have a significant effect on the abundance of piRNA pathway proteins. The chromatoid body (CB) is a large solitary electron dense structure devoted to RNA regulation in round spermatids^53^ and MVH/DDX4, a piRNA pathway protein, is a marker of the CB. Comparable CB structures, MVH expression and localization patterns in *Chr18^Δ/Δ^* and control mice are strongly supported by our transcriptome and proteome analyses (Extended Data Fig. 6b, c). Thus, *Chr18* piRNAs only affects post-transcriptional silencing during the final steps of differentiation in mouse spermiogenesis.

Using a luciferase assay system, we investigated the ability of the aforementioned annotated *Chr18* piRNAs to target cognate sequences of the 3’-UTR of *Golga2*. In agreement with a recent report^54^, we found that more than one *Chr18* piRNA can target the same cleavage site with a mismatch tolerance between n16 to 31 (Fig. 6b, bottom). Therefore, we constructed a mutant luciferase reporter with the deletion of the n2 to 31 sequence that includes both the piRNA seed sequence and the adjacent mismatched region. (Fig. 6c). To utilize endogenous endonuclease activity, GC-2spd(ts) cells which express MIWI^16^ were transfected with the luciferase reporter system. We found that both deletion of *Chr18* piRNA-binding site in the reporter and mutations in the corresponding piRNA completely abolished the repression effect of piRNA with detectable impact on reporter mRNA levels (Fig. 6d). This supports a direct mRNA target cleavage mechanism of *Chr18* piRNA through piRNA:mRNA base pairing. Together, it indicates that an accurate, post-transcriptional RNA regulatory mechanism regulated by *Chr18* piRNAs is critical for normal spermatogenesis.

Of note, other selected transcripts involved in acrosomal vesicle trafficking and fusion also were significantly increased in *Chr18^Δ/Δ^* testes compared with *Chr18^+/+^*and *Chr18^+/Δ^* controls (Fig. 6a). However, they did not have 3’-UTR piRNA seed sequences corresponding to the annotated *Chr18* piRNAs, suggesting that other mechanisms may pertain.

## Discussion

We have previously reported that BTBD18, a nuclear protein expressed in male germ cells, acts as a licensing factor required for transcriptional elongation of precursor transcripts at 50 of the 115 pachytene piRNA clusters present in the mouse genome. *Btbd18^Null^* spermatids undergo meiosis and arrest in the Golgi phase of spermiogenesis (stages 1-3). Elongated spermatids are not observed, and male germ cells undergo apoptosis^20^. We anticipated that ablation of individual binding sites of BTBD18 in the mouse genome would phenocopy a subset of *Btbd18^Null^* abnormalities. Indeed, disruption of the single pachytene piRNA producing locus on Chr18 resulted in notable phenotypic defects and male sterility. Thus, the severe defects observed in *Chr18^Δ/Δ^* mice appear to reflect a lower degree of functional redundancy with other piRNA clusters.

The relatively late expression of pachytene piRNA suggests a major role in regulating post-meiotic spermiogenesis^16, 17^. Despite *Miwi^Null^* mice failure to generate pachytene piRNA, round spermatids are generated, which indicates that meiosis can be completed without MIWI and MIWI-binding pachytene piRNAs^55, 56^. Instead, the main defect of *Miwi^Null^* mice is failure of haploid sperm differentiation that results in the absence of mature spermatozoa. Unlike *Miwi^Null^* and *Btbd18^Null^* mice, *Chr18^Δ/Δ^* mice can produce mature spermatozoa but are infertile. Consistent with predictions, the molecular defects of *Chr18^Δ/Δ^* mice occur during spermiogenesis in the transformation of haploid round spermatids to mature, elongated spermatozoa. Spermatogenesis was normal in *Chr18^Δ/Δ^* mice until the onset of acrosome biogenesis at the Golgi phase of spermiogenesis when sperm dysmorphology was associated with abnormalities in transcript and protein abundance.

The Chr18 piRNA cluster is ranked as the highest piRNA-producing locus among all (>200) pre- and pachytene piRNA clusters^28^. While *Chr18^+/Δ^* control mice had normal fertility and no discernable phenotype, transcriptome analysis revealed minor changes of mRNA abundance between *Chr18^+/Δ^* and *Chr18^Δ/Δ^* testes which suggests compensatory mechanisms that could partially account for the difficulty in identifying pachytene piRNA targets and function^57, 58^. Ablation of the promoter of the single bi-directional site on Chr18 causes a phenotype that develops later in spermiogenesis compared to *Btbd18^Null^* mice. *Chr18^Δ/Δ^* spermatids can elongate albeit with severe head dysmorphology that affects acrosome reaction and poor progressive motility that render male mice infertile. Comparison of the transcriptomes and proteomes of *Chr18^Δ/Δ^* testes with *Chr18^+/+^*and *Chr18^+/Δ^* controls suggests that removal of a subset of piRNAs can increase mRNA abundance which resulted in disruption of protein homeostasis, but whether this is mediated transcriptionally or post-transcriptionally (or both) remains unclear. However, a recent observation proposed that a subset of pachytene piRNAs can regulate target mRNA translation through seed complementarity in their target mRNAs 3’-UTR without alteration on mRNA levels^59^.

The primary alteration in spermiogenesis in *Chr18^Δ/Δ^* mice was the loss of structural integrity of the trans-Golgi network (TGN) which appeared as loose whorls associated with enlarged proacrosomal vesicles (PV) compared to controls. The loss of Chr18 pachytene piRNA induced enhanced PV formation and trafficking that resulted in dramatic acrosomal overgrowth. Although post-transcriptional repression of mRNA targets by miRNAs is well documented^60^, a potential role for pachytene piRNAs in targeting specific RNAs for degradation also has been reported^16, 17, 54^. Chr18 piRNAs can bind with complementarity to *Golga2* transcripts by pairing to nt 2-15 of the piRNA seed and cleaving the target 3’-UTR^18^ which would lead to degradation of the transcript. Thus, the absence of the piRNA from the Chr18 cluster could account for the increased abundance of *Golga2* transcripts in the *Chr18^Δ/Δ^* mice.

Several gene-edited mouse models, including *Golga2^Null^* mice, with defects in acrosome biogenesis result in loss of acrosome formation and globozoospermia which is a phenotype seen in infertile humans^33, 44, 61–68^. During the Golgi phase of spermiogenesis in *Smap2^Null^* mice (lacking an arf GTPase-activating protein), spermatids had similar defects of PV formation and distorted TGN structure and yet produced globozoospermia^64^. In addition, in the absence of zona pellucida binding protein 2, *Zpbp2^Null^* spermatozoa have subtle head deformations with shortened apical hooks and bulges but were still able to undergo acrosome exocytosis^63^. Moreover, in the absence of proprotein convertase 4, *Pcsk4^Null^* mice had a sickle-shaped head and lacked the pointed apex similar with *Chr18^Δ/Δ^* sperm. However, mRNA abundance and protein level of PCSK and its substrate, acrosin-binding protein, ACRBP were not altered in *Chr18^Δ/Δ^* mice^65^. Therefore, the observed acrosomal overgrowth and impaired acrosome exocytosis appear unique to *Chr18^Δ/Δ^* mice and reflect the functional significance of Chr18 pachytene piRNA as a regulator for acrosome biogenesis during the spermiogenesis.

Except for *Golga2*, the altered proteomic profiles do not correlate with corresponding changes in the transcriptome of *Chr18^Δ/Δ^* mice. However, the increased level of HSPs and UPS in *Chr18^Δ/Δ^* mice suggests that the absence of Chr18 pachytene piRNAs affects quality monitoring systems involved folding, refolding and degradation of proteins. RNA-seq documented that mRNA abundance of *Odf2*, *Bag5*, *Stub1*, *Boll*, and *Hspa2* were identical in *Chr18^Δ/Δ^* and control mice. When protein quality control systems fail, damaged proteins accumulate as aggregates. HSPs (renaturation) act as molecular chaperones and the UPS (degradation) compete in the selective degradation of damaged or abnormal proteins. While it was not originally included in our screen for proteins involved in protein triage, TIA-1 is increased in *Chr18^Δ/Δ^* mice. It has been shown that the main function of stress proteins is to prevent accumulation of denatured and/or aggregated proteins, as an increase in ubiquitin-dependent degradation of proteins could present heat shock toxicity as efficiently as HSPs^69^. In addition, our proteomic analysis identified increased amounts of stress protein and eukaryotic translation initiation factor 3 (eiF3) which is required for stress granule formation in mammals^70^.

In summary, we have used *Chr18^Δ/Δ^* mice to investigate molecular mechanisms by which pachytene piRNAs can regulate mRNA abundance and triage proteins to ensure production of functional spermatozoa and successful fertilization. There are continuous compensatory mechanisms to overcome failure to maintain mRNA and protein homeostasis which may account for the modest molecular perturbations observed by RNA-seq and microscale mass spectrometry analyses. Moreover, imbalance between HSPs (renaturing) and UPS (destroying) leads to high sensitivity to stress. These results suggest a previously unrecognized relationship between protein triage, stress response, and pachytene piRNAs. Thus, the *Chr18^Δ/Δ^* gene-edited mouse model may provide insight into the role of pachytene piRNAs beyond the dramatic effect on spermiogenesis.

## METHODS

### Animals

All experiments with mice were conducted in accordance with guidelines of the National Institute of Health under a Division of Intramural Research and NIDDK Animal Care and Use Committee approved animal study protocol (protocol numbers KO18-LCDB-18 and KO44-LCDB-19).

### Generation of CRISPR/Cas9 mutant mice

To establish *Chr18^Δ/Δ^* mutant mice, the single guide RNA (sgRNA) sequences (Extended Data Fig. 1c) were designed to target the bi-directional promoter and flanking sequence of the Chr18 piRNA cluster. Synthetic double-stranded DNA was cloned into pDR274 (Addgene, #42250) to express a sgRNA. After digestion with *Dra*I, the linearized DNA fragment was purified with a PCR Clean-up Kit (Clontech Laboratories) and *in vitro* transcribed using the AmpliScribe T7-Flash Transcription Kit (Lucigen). *Cas9* cRNA (Addgene #42251) was generated after linearization with *Pme*I, purified with the PCR clean-up kit, and *in vitro* transcribed with mMESSAGE mMACHINE T7 (Thermo Fisher Scientific). Both sgRNA and *Cas9* cRNA were purified with MEGAclear Transcription Clean-Up Kit (Thermo Fisher Scientific). To collect zygotes from oviducts at embryonic day 0.5 (E0.5), hormonally stimulated B6D2_F1_ (C57LB/6 × DBA2) female mice were mated with B6D2_F1_ male mice. Mixed sgRNA (50 ng/μl) and *Cas9* cRNA (100 ng/μl) were injected into zygotes in M2 medium. Injected zygotes were cultured (12-18 hr) in KSOM (37 °C, 5% CO_2_) supplemented with 3 mg/ml BSA to two-cell embryos and transferred into oviducts of pseudo-pregnant ICR female mice. To determine the genotype of mutant founders, genomic DNA was extracted from tail tips and lysed in 150 μl of DirectPCR Lysis Reagent (Viagen Biotech) with protease K (0.2 mg/ml, Thermo Fisher Scientific) at 55 °C for 5 hr. After protease K inactivation by incubation at 85 °C for 1 hr, samples were genotyped by PCR. After purification, PCR products were cloned into TOPO blunt vectors for DNA sequencing. Mouse mutant lines were established and maintained by mating mutant founders with C57BL/6 females or males. All mutant mice in this study were backcrossed for at least two generations before use.

### Mouse sperm preparation

Sperm from cauda epididymides were released into Cook medium (Cook Medical) and squeezed from vas deferens.

### Fertility

To assess fertility, individual 2-8 mo old male mice were co-caged with two C57LB/6 females for 2 wk to 6 mo. The average number of pups per litter was quantified and at least 5 mating cages were set up for each genotype. Female mice were checked for the presence of vaginal plugs and pregnancy. The same procedures were used to assess the fertility of *Chr18^Δ/Δ^* and control female mice.

### *In vitro* fertilization

To assess *in vitro* fertility, caudal epididymal sperm were isolated from 2-8 mo old *Chr18^Δ/Δ^* and control mice and capacitated for 1.5 to 2 hr in 0.5 ml of Cook medium (Cook Medical). Wild-type B6D2_F1_ female mice (2-3 mo old) were synchronized with 5 U of PMSG and induced to ovulate with 5 U of hCG administered 48 hr later. Cumulus-intact eggs were recovered from oviducts 15 to 16 hr later in 0.2 ml of Cook medium with 1 mM reduced glutathione (GSH, Sigma). To minimize differences in the quality of recovered eggs, cumulus-intact eggs in one oviduct were separated from those in the other oviduct. Capacitated sperm (1.5 × 10^5^/ml) were added to each pool and incubated for 3-6 hr (37 °C, 5% CO_2_ in air). The eggs were transferred to KSOM (37 °C, 5% CO_2_) supplemented with 4 mg/ml BSA and the presence of two pronuclei was recorded as fertilized.

### Sperm count, motility, and morphology analysis

To count sperm, cauda epididymides and vas deferens were harvested in pre-warmed (37 °C) Cook medium (Cook Medical). 20 μl of a sperm suspension was diluted in 500 μl of Cook medium and counted in a hemocytometer using an AxioPlan 2 (Carl Zeiss) microscope. Isolated sperm motility was determined by computer assisted sperm analysis (CASA) using a HTM-IVOS (Version 12.3) motility analyzer (Hamilton Thorne). Sperm were further observed for morphological changes by light microscopy after staining for DNA with Hoechst 33342. In addition, acrosome exocytosis of the isolated sperm was induced by 20 μM calcium ionophore (A23187, Sigma Aldrich) in pre-warmed HTF media followed by incubation (37 °C, 5% CO_2_, 90 min).

### Isolation of primary germ cells

Testes were excised to remove seminiferous tubules which were minced gently with fine scissors over 3–4 min in 1 ml of M16 medium (Sigma Aldrich). The minced tissue was treated with 0.05% collagenase/trypsin^71^. The resultant cell suspension was washed with and resuspended in M2 medium (Sigma Aldrich).

### Histology, TUNEL, immunofluorescence and confocal microscopy

Mouse testes and epididymides were incubated in Bouin’s fixative overnight at room temperature (RT) and washed with 70% EtOH. Paraffin embedded samples were sectioned (5 μm) and mounted on slides prior to staining with periodic acid-Schiff (PAS) and hematoxylin. Stages of spermatogenesis and steps of spermatid development were determined^72^. Terminal deoxynucleotidyl transferase-mediated deoxyuridine triphosphate (TUNEL) assay was used to determine apoptosis with an *In Situ* Apoptosis Detection Kit (Millipore) according to the manufacturer’s instructions. Bright field images were obtained with an AxioPlan 2 (Carl Zeiss) microscope.

After deparaffinization, rehydration, and antigen retrieval with 0.01% sodium citrate buffer (pH 6.0) (Sigma Aldrich), tissue sections were incubated with blocking buffer (3% goat serum, 0.05% Tween-20, RT for 1 hr) followed by addition of primary antibodies (Supplementary Table 2) overnight at RT. Specific Alexa Fluor secondary antibodies were used to detect primary antibodies and DNA was stained with Hoechst 33342. Fluorescent images were captured with an LSM 780 (Carl Zeiss) confocal/multiphoton microscope.

### Meiotic chromosome spreads

To obtain meiotic chromosome spreads, de-capsuled mouse testes were incubated in hypotonic extraction buffer (30-60 min, on ice) and seminiferous tubules were chopped to release germ cells^73^. A drop of sucrose solution containing germ cells was placed on a glass slide coated with 1× PBS containing 1% PFA and 0.15% (v/v) Triton-X100 (pH 9.2) and spread by swirling. Slides were placed in a humidifying chamber (2.5 hr), air-dried, washed twice with 1× PBS with 0.4% Photo-Flo 200 solution (Electron Microscopy Science) and air-dried. For immunostaining of meiotic chromosomes, slides were blocked with blocking buffer (3% goat serum, 0.05% Tween-20, RT for 1 hr). The slides were then incubated with primary antibodies in a humidifying chamber overnight at RT. The slides were incubated with Alexa Fluor secondary antibodies (1 hr, RT). Hoechst was used for DNA staining and images were obtained with an LSM 780 confocal/multiphoton microscope.

### Isolation of primary germ cells

Testes were excised to remove seminiferous tubules which were minced gently with fine scissors (3–4 min) in 1 ml of M16 medium (Sigma Aldrich). The minced tissue was treated with 0.05% collagenase/trypsin^71^. The cell suspensions were washed with and resuspended in M2 medium (Sigma Aldrich). Cell suspensions were spread on poly-lysine-coated glass slides (15 min, RT) and then fixed in cold methanol (−20 °C, 15 min). The slides were dried for 10 min to evaporate methanol, treated with 0.2% Triton X-100 for 10 min followed by four washings with PBS. The slides were placed in blocking buffer (PBS containing 3% goat serum, 1% glycerol, 0.1% BSA) for 60 min at RT and incubated overnight with primary antibodies at RT. After washing with PBS twice for 10 min, samples were incubated with secondary Alexa Fluor conjugated antibody for 60 min. The slides were mounted using antifade mounting medium and images were obtained with an LSM 780 confocal/multiphoton microscope.

### Scanning and transmission electron microscopy

For scanning electron microscopy (SEM), sperm from cauda epididymides and vas deferens were isolated in 0.1 M phosphate buffer, pH 7.4. Sperm were attached to a poly-L-lysine coated glass coverslip and fixed (2.5% glutaraldehyde, 1% formaldehyde, 0.12 M sodium cacodylate buffer, pH 7.4, 1 hr, RT). Samples were washed in cacodylate buffer, post-fixed (1% osmium tetroxide, 1 hr) dehydrated in a graded ethanol series, and dried out of CO_2_ in a Samdri-795 critical point dryer (Tousimis Research Corp). Samples were mounted on SEM stubs with carbon adhesive, sputter-coated with 5-10 nm of gold in an EMS 575-X sputter coater (Electron Microscopy Sciences) and examined with a ZEISS Crossbeam 540 SEM at the NHLBI Electron Microscopy Core.

For transmission electron microscopy (TEM), testes, cauda epididymides, and vas deferens sperm were fixed (2.5% glutaraldehyde, 1% formaldehyde, 0.12 M sodium cacodylate buffer, pH 7.3), cut into 1 mm^3^ pieces, post-fixed (1% osmium tetroxide) and stained (1% uranyl acetate). Samples were dehydrated and embedded in Epon 812 resin. Ultrathin sections were counterstained with uranyl acetate and lead citrate. Images were acquired with a JEM 1200EX TEM equipped with an AMT 6-megapixel digital camera at the NHLBI Electron Microscopy Core. Densitometric quantification of acrosomal granules and acrosomal vesicles and diameter of nuclei were processed with ImageJ software (NIH).

### Protein extraction and immunoblots

Testicular and sperm proteins were extracted in 1× LDS sample buffer with 1× NuPAGE Sample Reducing Agent (Thermo Fisher Scientific). Proteins were separated on 4-12% Bis-Tris gels and electrophoretically transferred to PVDF membranes. The membranes were blocked with 5% nonfat milk in Tris-buffered saline containing 0.05% Tween-20 (TBS-T) at RT for 1 hr and probed with primary antibodies (Supplementary Table 2) overnight at 4 °C. The membranes were washed three times with TBS-T and incubated for 1 hr at RT with secondary antibodies followed by washing with TBS-T and developed using SuperSignal West Dura Extended Duration Substrate (Thermo Fisher Scientific). Signals were detected with PXi Touch (Syngene) according to the manufacturer’s instructions. For immunoblot analysis, sperm from the cauda epididymides and vas deferens were directly released into PBS. The collected sperm were washed with PBS and then resuspended in sample buffer containing 3% SDS and boiled for 10 min.

### RNA-seq library preparation

Total RNA (100-1000 ng) was isolated from testes using miRNAeasy Mini (Qiagen) and RNA-seq libraries were constructed using TruSeq Stranded Total RNA Kit (Illumina) with Ribo-Zero following the manufacturers’ instruction. The fragment size of RNA-seq libraries was verified using a 2100 Bioanalyzer (Agilent) and the concentrations were determined using Qubit (LifeTech). The libraries were loaded onto the Illumina HiSeq 3000 for 2×50 bp paired end read sequencing at the NHLBI DNA Sequencing and Genomics Core Facility. The fastq files were generated using the bcl2fastq software for further analysis.

### Small RNA-seq library preparation

As previously described for preparation of small RNA-seq libraries^74^, total testes RNA (20 µg) was incubated with 4 μl of 5X borate buffer (148 mM borax, 148 mM boric acid, pH 8.6, Thermo Fisher Scientific) for 10 min at RT with 2.5 μl of freshly dissolved 200 mM NaIO_4_ (Thermo Fisher Scientific) for β-elimination. To quench unreacted NaIO_4_, 2 μl of glycerol (ThermoFisher Scientific) was added and incubated for 10 min at RT. After adding 380 μl of 1X borate buffer, RNA was precipitated with ethanol for 1 hr, at −80 °C. Following centrifugation, the RNA was dissolved in 50 μl of 1X borax buffer (30 mM borax and 30mM boric acid, 17.5 mM NaOH, pH 9.5) and incubated for 90 min at 45 °C prior to addition of 450 μl of 1X borate buffer and 20 μg of glycogen. The RNA was precipitated with ethanol for 1 hr, at -80 °C, collected by centrifugation and dissolved in water. During β-elimination, periodate-reacted RNAs were shortened by 1 nt at the 3’ end with monophosphates and were unable to be amplified during library preparation. Thus, piRNAs, protected from β-elimination by 2’-O-methylation at the 3’ end, were enriched in the small RNA-seq libraries. For small RNA-seq library construction, NEBNext Multiplex Small RNA Library Prep Set for Illumina (New England BioLabs) was used per the manufacturer’s instructions. In general, 1 μg total RNA was subjected to 3’ and 5’ adapter ligation, reverse transcribed, PCR amplified, followed by size selection with AMPure XP beads (Beckman Coulter) for deep sequencing at the NIDDK Genomics Core Facility.

### RNA-seq data analysis

Raw sequence reads were trimmed with cutadapt 1.18 to remove any adapters while performing light quality trimming with parameters “-a ATCGGAAGAGC -A ATCGGAAGAGC -q 20 --minimum-length=25.” Sequencing library quality was assessed with fastqc v0.11.8 with default parameters^75^ and trimmed reads were mapped to the *Mus musculus* mm10 reference genome using hisat2 2.1.0 with default parameters^75^. Multimapping reads were filtered using SAMtools 1.9^76, 77^. Uniquely aligned reads were then mapped to gene features using subread featureCounts v1.6.2 as a second strand library with parameters^78^. “-t gene -g gene_id -f -p -B -P -C.” Differential expression between groups of samples was tested using R version 3.5.1 (2018-07-02) with DESeq2 1.20.0^79^. Transcript quantification was performed with salmon 0.11.3 with parameters^80^ “--gcBias --libType A --seqBias --threads 1.” piRNA annotations were derived from the Zamore lab^3, 81^. Transposon-mapping reads were aligned to repBase annotated regions^82^, upbuilt from mm9 to mm10, using the software pipeline piPipes and bowtie2 2.2.5^83^.

### Small RNA-seq data analysis

After removing adaptors, rRNA and tRNA sequences were filtered. The remaining reads with sizes from 26 to 31 nt were mapped to the UCSC mm 10 assembly using hisat2 2.1.0^75^ and only uniquely mapped reads were used for further analysis. Through miRNA counts (miRbase) normalization piRNA abundance was obtained.

### Quantitative real-time RT-PCR (qRT-PCR)

Total RNA was isolated from mouse tissues using a miRNAeasy Mini Kit (Qiagen) and cDNA was synthesized with a RevertAid Premium First Strand cDNA Synthesis Kit (Thermo Fisher Scientific). Quantitative RT-PCR was performed using iTaq Universal SYBR Green Supermix (Bio-Rad) and QuantStudio 6 Flex Real-Time PCR System (Thermo Fisher Scientific). The relative abundance of each transcript was calculated by the 2^−ΔΔ*Ct*^ normalized to endogenous *β-actin* expression^84^.

### Tandem Mass Tag (TMT) mass spectrometry

Mass spectrometry was performed at the Harvard FAS Division of Science Mass Spectrometry and Proteomics Resource Laboratory.

#### Lysis, digestion and labeling procedures

All cells were placed into Covaris^®^ microTUBE-15 (Woburn) microtubes with Covaris^®^ TPP buffer. Samples were lysed in Covaris S220 Focused-ultrasonicator with 125W power over 180 sec with 10% max peak power. Lysed cells were chloroform/MeOH precipitated, weighed and then digested via FASP digest as described: new 10 K filter were washed with 100 μl of 1 M Triethylammonium bicarbonate (TEAB) prior to adding 100 μl of 0.5 M Tris-(2 carboxyethyl) phosphine (TCEP) and incubating at 37 °C for 1 hr. IAcNH_2_ (100 μl) was added in the dark for 1 hr and incubated with 150 μl of TEAB and trypsin (Promega) overnight at 38 °C. 50 mM TEAB (50 μl) was added prior to HPLC.

#### TMT Mass tagging and peptide labeling

Immediately before use, the TMT Label Reagents (Thermo Fisher Scientific) were equilibrated to RT. For the 0.8 mg vials, 41 μl of anhydrous acetonitrile or ethanol was added to each tube. For the 5 mg vials, 256 μl of solvent was added to each tube. The reagents were allowed to dissolve for 5 min with occasional vortexing. The tube was briefly centrifuged to gather the solution and 41 μl of the TMT Label Reagent was carefully added to each 25-100 μg sample. Alternatively, the reduced and alkylated protein was transferred to the TMT Label Reagent vial and incubated for 1 hr at RT. 8 μl of 5% hydroxylamine to was added to the sample and incubated for 15 min to quench the reaction. Samples were combined in equal amounts and stored at -80 °C. The pooled samples were divided into 20 fractions and submitted for LC-MS/MS performed on an Orbitrap Lumos (Thermo Fisher Scientific) equipped with EASYLC1000 (Thermo Fisher Scientific). Peptides were separated onto a 100 µm inner diameter microcapillary trapping column packed first with approximately 5 cm of C18 Reprosil resin (5 µm, 100 Å, Dr. Maisch GmbH) followed by an analytical column ∼20 cm of Reprosil resin (1.8 µm, 200 Å, Dr. Maisch GmbH). Separation was achieved by applying a gradient from 5-27% acetonitrile in 0.1% formic acid over 90 min at 200 nl/min. Electrospray ionization was enabled by applying a voltage of 1.8 kV using a home-made electrode junction at the end of the microcapillary column and sprayed from fused silica pico tips (New Objective). The LTQ Orbitrap Lumos was operated in data-dependent mode for mass spectrometry. The mass spectrometry survey scan was performed in the Orbitrap in the range of 395-1,800 m/z at a resolution of 6 × 10^4^, followed by the selection of the twenty most intense ions (TOP20) for CID-MS2 fragmentation in the Ion trap using a precursor isolation width window of 2 m/z, AGC setting of 10,000, and a maximum ion accumulation of 200 ms. Singly charged ion species were not subjected to CID fragmentation. Normalized collision energy was set to 35 V and an activation time of 10 ms. Ions in 10 ppm m/z window around ions selected for MS2 were excluded from further selection for fragmentation for 60 s. The same TOP20 ions were subjected to HCD MS2 event in the Orbitrap part of the instrument. The fragment ion isolation width was set to 0.7 m/z, AGC was set to 50,000, the maximum ion time was 200 ms, normalized collision energy was set to 27V and an activation time of 1 ms for each HCD MS2 scan.

#### Mass spectrometry analysis

Raw data were submitted for analysis in Proteome Discover 2.2 (Thermo Fisher Scientific) software. Assignment of MS/MS spectra was performed using the Sequest HT algorithm by searching the data against a protein sequence database including all entries from the Uniprot_Mouse2018_SPonly.fasta database as well as other known contaminants such as human keratins and common lab contaminants. Sequest HT searches were performed using 20 ppm precursor ion tolerance and requiring each peptides N and C termini to adhere with trypsin protease specificity, while allowing up to two missed cleavages. 6-plex TMT tags on peptide N termini and lysine residues (+229.162932 Da) was set as static modifications while methionine oxidation (+15.99492 Da) was set as variable modification. A MS2 spectra assignment false discovery rate (FDR) of 1% on both protein and peptide level was achieved by applying the target-decoy database search. Filtering was performed using a Percolator (64bit version). For quantification, a 0.02 m/z window centered on the theoretical m/z value of each the six reporter ions and the intensity of the signal closest to the theoretical m/z value was recorded. The listed proteins of significantly changed with a *P* < 0.05 was used to generate volcano plots and for Qiagen’s Ingenuity Pathway Analysis (IPA, Qiagen).

### Plasmids and RNA oligonucleotides

For reporter pRL-*Golga2* 3’-UTR, the ∼1.3 kb mouse *Golga2* 3’-UTR was cloned downstream of the *Renilla* luciferase gene in pRL-TK (Promega). Thirty-one nucleotides in *Golga2* 3’-UTR complementary to the 5’ portion of *Chr18* piRNA were deleted for generating reporter *Golga2* 3’-UTR Mut by Gibson assembly (New England Biolabs). All constructs were confirmed by DNA sequencing. All piRNA oligonucleotides were synthesized by IDT and sequences are provided in supplementary information. All chemically synthetic piRNAs used for cell culture transfection are 2’-O-methyl modified at their 3’ ends.

### Cell culture and transfection

GC-2spd(ts) cells were obtained from the American Type Culture Collection (ATCC) and cultured with the medium and serum as ATCC recommended. Transfection was performed using Lipofectamine 3000 (Invitrogen) according to the manufacturer’s instructions. For transfection of the RNA oligonucleotides, 500 nM of piRNA oligonucleotides were used.

### Dual luciferase reporter assay

Dual luciferase reporter assays were carried out as described^16^. In brief, GC-2spd (ts) cells were co-transfected in 96-well plates with chemically synthetic piRNAs and *Renilla* luciferase 3’-UTR reporters. The firefly luciferase plasmid pGL4 was used as an internal control. Dual luciferase assays were performed 24 h after transfection using Dual-Glo Luciferase Assay Systems (Promega).

### Quantification and statistical analysis

All statistical analyses were performed using Graph Pad Prism 8 software. For details of the detailed statistical analyses used, precise *P* values, statistical significance, and sample sizes for all of the graphs, see the figure legends. No statistical methods were used to predetermine sample size, experiments were not randomized, and investigators were not blinded to allocation during experiments and outcome assessment, unless stated otherwise.

## Data availability

The accession number for the sequencing data reported in this paper is GEO (Gene Expression Omnibus): GSE136603. The datasets generated during the current study are available from the corresponding authors upon reasonable request.

## Acknowledgements

We thank members of the Dean lab for helpful suggestions, Dr. Liquan Zhou for initial advice on targeting strategies to establish mutant mouse line, and Dr. Cameron D. Palmer for bioinformatic assistance. We appreciate the critical reading of the manuscript by Drs. Astrid D. Haase and P. Jeremy Wang. We thank the NIDDK and NHLBI Genomics and Electron Microscopy Core for deep sequencing and SEM/TEM analyses. We also thank the Harvard University FAS Division of Science MSPRL for proteomic analysis. This work was supported by the Division of Intramural Research Program of the National Institutes of Health, NIDDK.

## Author contributions

H.C. and J.D. designed the experiments, analyzed the data, and wrote the paper. H.C. performed the experiments. Z.W. performed mouse embryo injections.

## Extended Data

**Extended Data Fig. 1.**
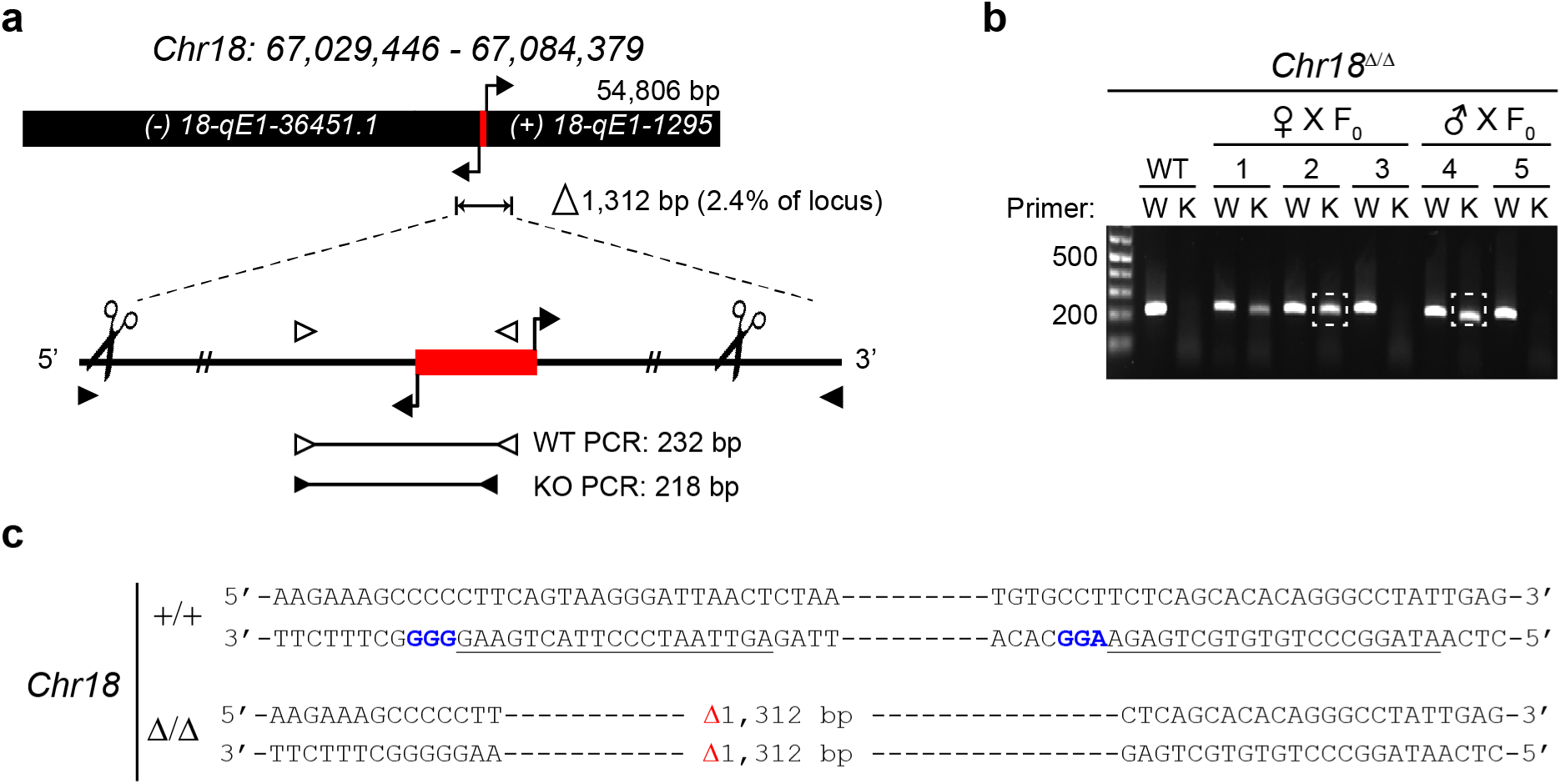
Generation of *Chr18^Δ/Δ^* mice. **a**, Schematic diagram of Chr18 piRNA coding locus deleted using CRISPR/Cas9. Scissors, sgRNAs target sites used to guide the Cas9-catalyzed promoter; red boxes, deletion. **b**, Genotyping of mutant founders by PCR. **c**, Genomic sequences of Chr18 piRNA promoter region in *Chr18^Δ/Δ^*. Dashes, genomic sequences deleted by CRISPR; blue NGG is protospacer adjacent motif (PAM); underlined, sgRNA.

**Extended Data Fig. 2.**
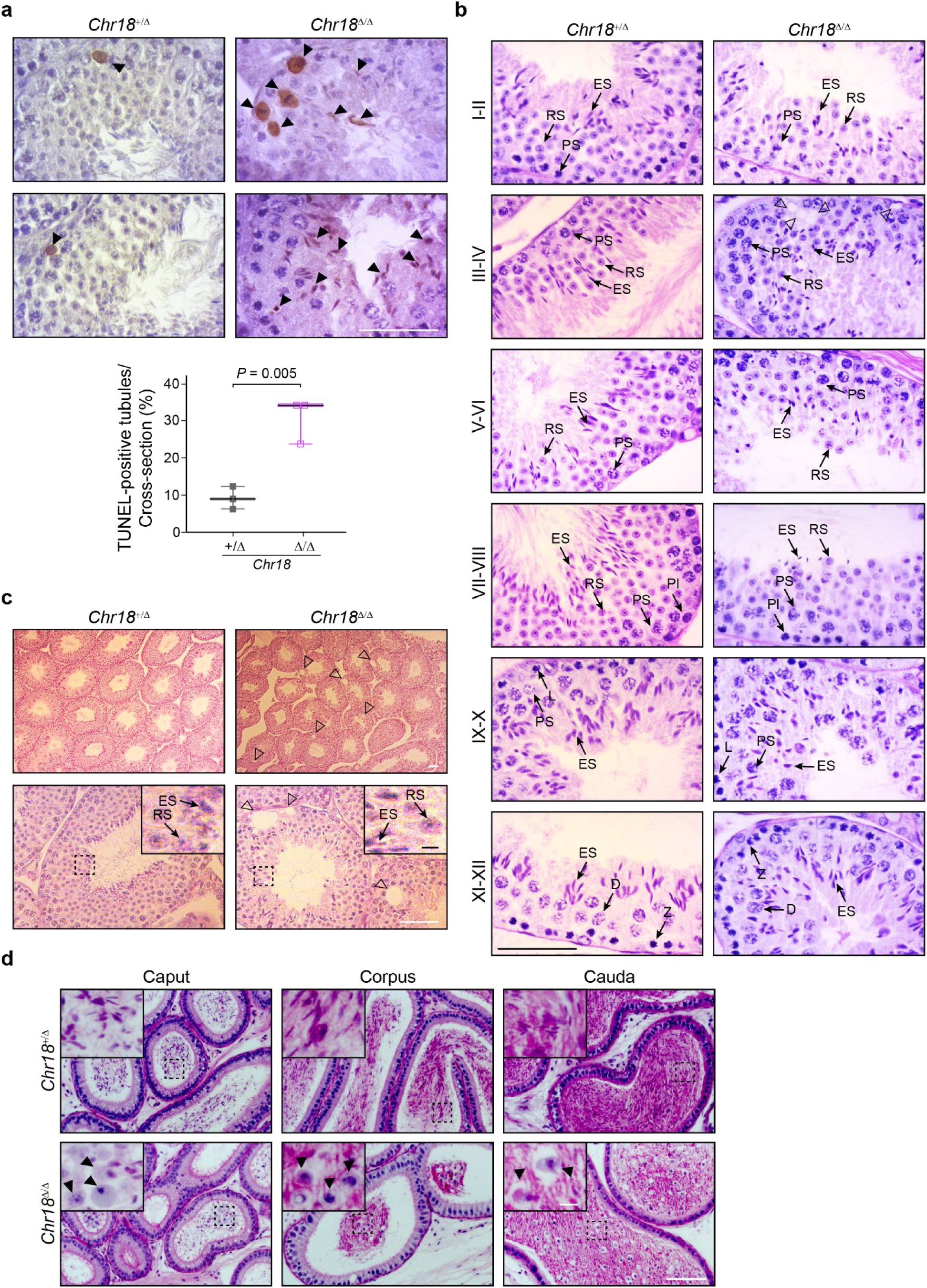
Histological defects and increased apoptosis in *Chr18^Δ/Δ^* mice. **a**, TUNEL staining (top) and quantification (bottom) of TUNEL-positive tubules per cross-section from 8 wk/old mice (*n* = 3). Black arrow heads, apoptotic cells. Scale bar, 50 μm. The box indicates median ± interquartile range, the whiskers indicate the highest/lowest values and midlines are median values. Two-tailed *P* values were calculated using Student’s *t*-test. **b**, Testicular sections from 8 wk/old mice were stained with periodic acid-Schiff (PAS) and hematoxylin (H) to determine stages of seminiferous epithelium cycles (*n* = 3). Pl, preleptotene; L, leptotene; Z, zygotene; PS, pachytene; D, diplotene; RS, round spermatids; ES, elongating spermatids. Scale bar, 50 μm. Stage of seminiferous epithelium cycles was determined by morphology of spermatocytes and round spermatids stained with PAS. **c**, Representative light microscopic images of testicular sections from 8 wk/old mice (n = 3 for each genotype). RS, round spermatid; ES, elongating spermatid; arrow heads, vacuolation. Scale bar, 50 μm; inset scale bar, 5 μm. **d**, PAS&H staining of epididymides from 12 wk/old mice (*n* = 2). Black arrow heads, sloughing germ cells. Scale bar, 50 μm; inset scale bar, 5 μm. **c**, **d** The insets are enlargements of the dashed box regions.

**Extended Data Fig. 3.**
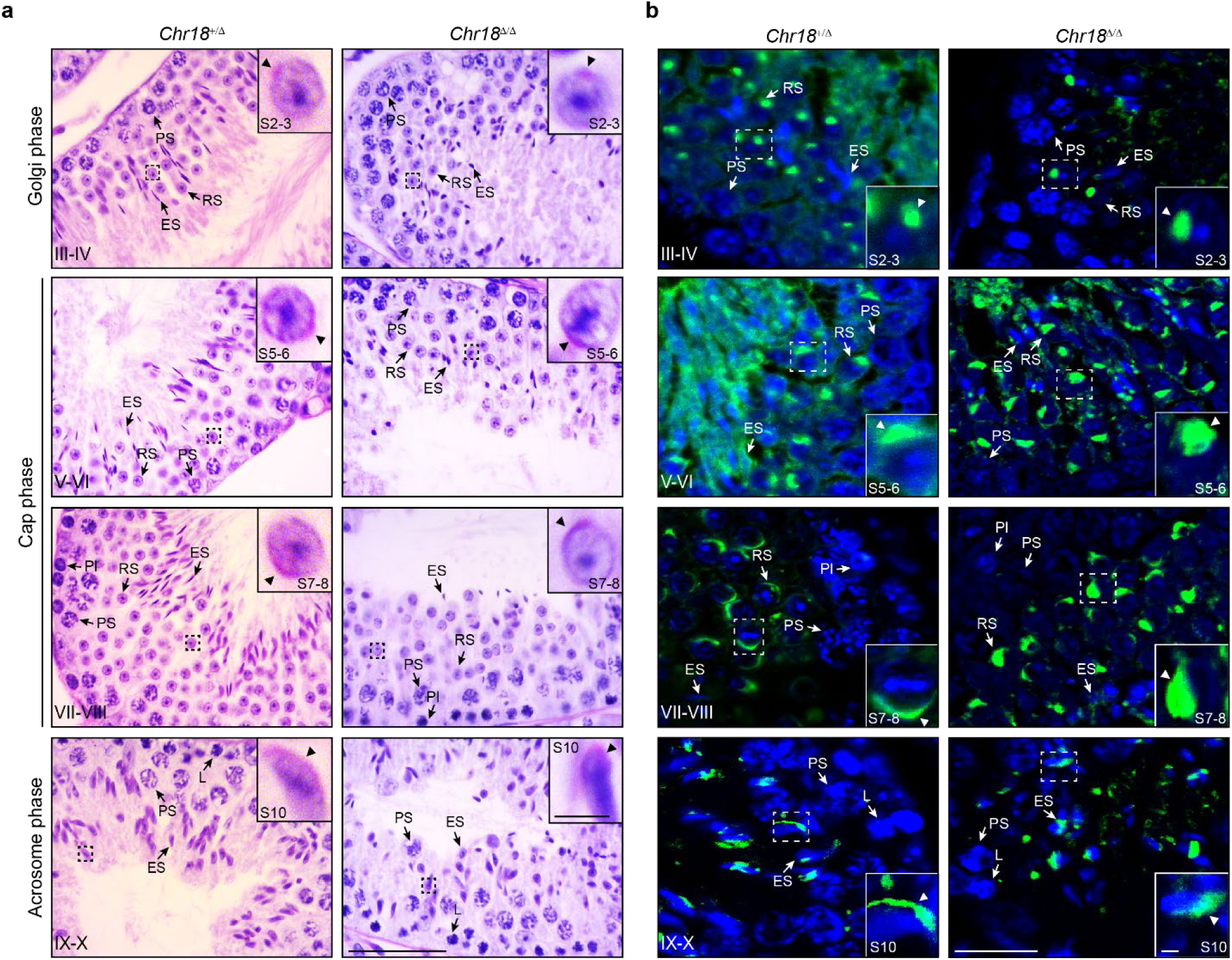
Acrosomal defects in *Chr18^Δ/Δ^* spermatids. **a**, **b**, Representative light microscopic images of PAS&H (**a**) and PNA staining (**b**) of testicular sections to evaluate acrosome biogenesis: Golgi, cap, and acrosome phases from 8 wk/old control and mutant mice (n = 3). The insets are enlargements of the dashed box regions. Pl, preleptotene; PS, pachytene spermatocyte; RS, round spermatid; ES, elongating spermatid; arrow heads, acrosome. Scale bar, 50 μm; inset scale bar, 5 μm. Sections were stained with PNA (acrosome, green) and Hoechst 33342 (DNA, blue). S2-8, step 2-8 round spermatids; S10, step 10 elongating or elongated spermatids. Stage of seminiferous epithelium cycles was determined by morphology of spermatocytes and round spermatids stained with PAS (**a**) and PNA (**b**). **a**, **b** The insets are enlargements of the dashed box regions.

**Extended Data Fig. 4.**
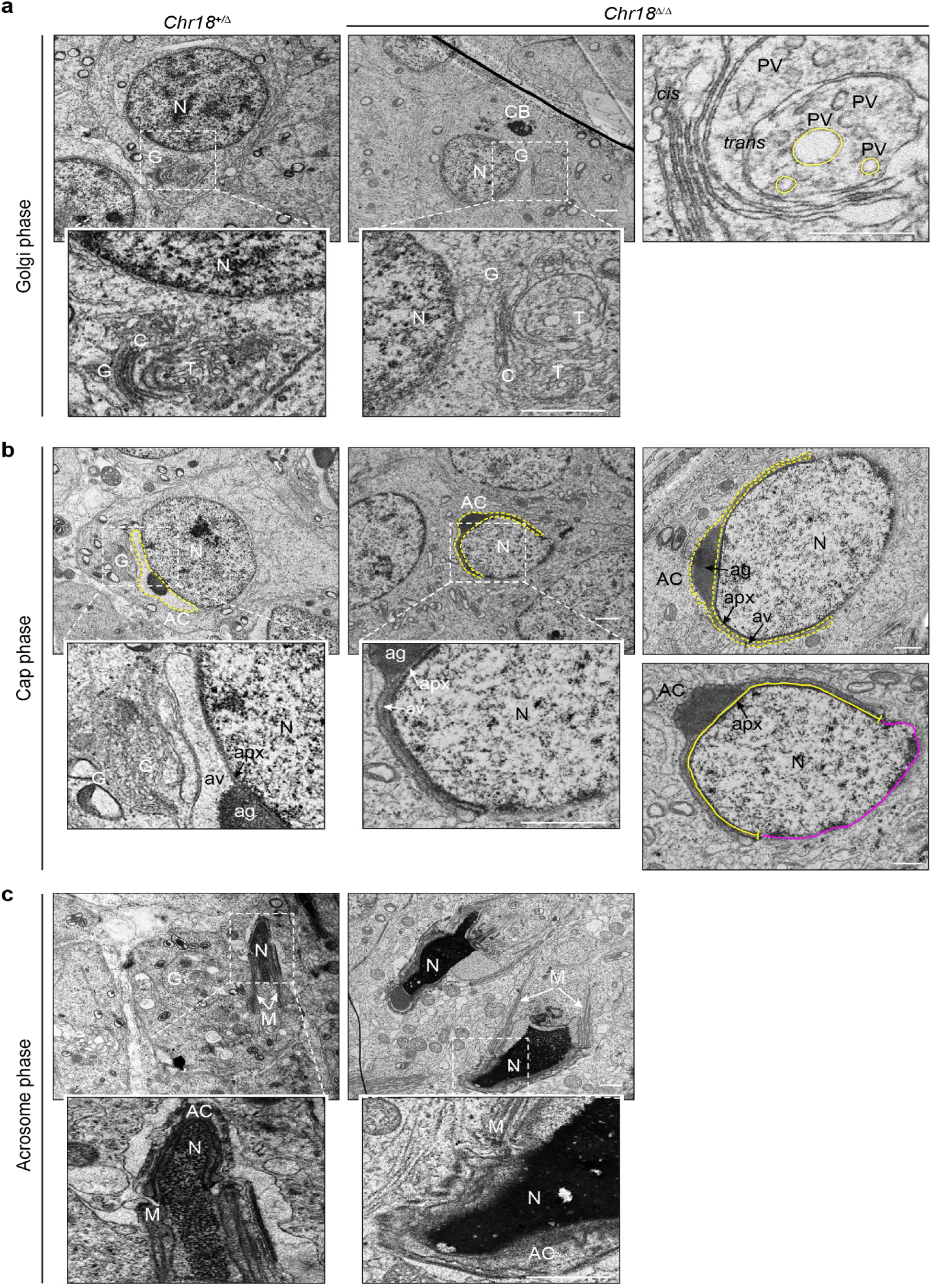
Acrosomal defects in *Chr18^Δ/Δ^* spermatid. **a-c**, Representative TEM images of Golgi (**a**), cap (**b**), and acrosome (**c**) phase spermatids from 12 wk/old control and mutant mice (*n* = 2 for each genotype). N, nucleus; G, Golgi apparatus; C, *cis*-Golgi network; T, *trans*-Golgi network; PV, proacrosomal vesicle; CB, chromatoid body; AC, acrosome; Ag, acrosomal granule; Av, acrosomal vesicle; Apx, acroplaxome; M, manchette. Scale bar, 0.5 μm. ImageJ was used to quantify the density of ag (black area) and av (yellow dashed area) (top) and measure the nuclear perimeter (yellow and purple area) and length of apx (yellow line) (bottom) in Fig. 4 of manuscript.

**Extended Data Fig. 5.**
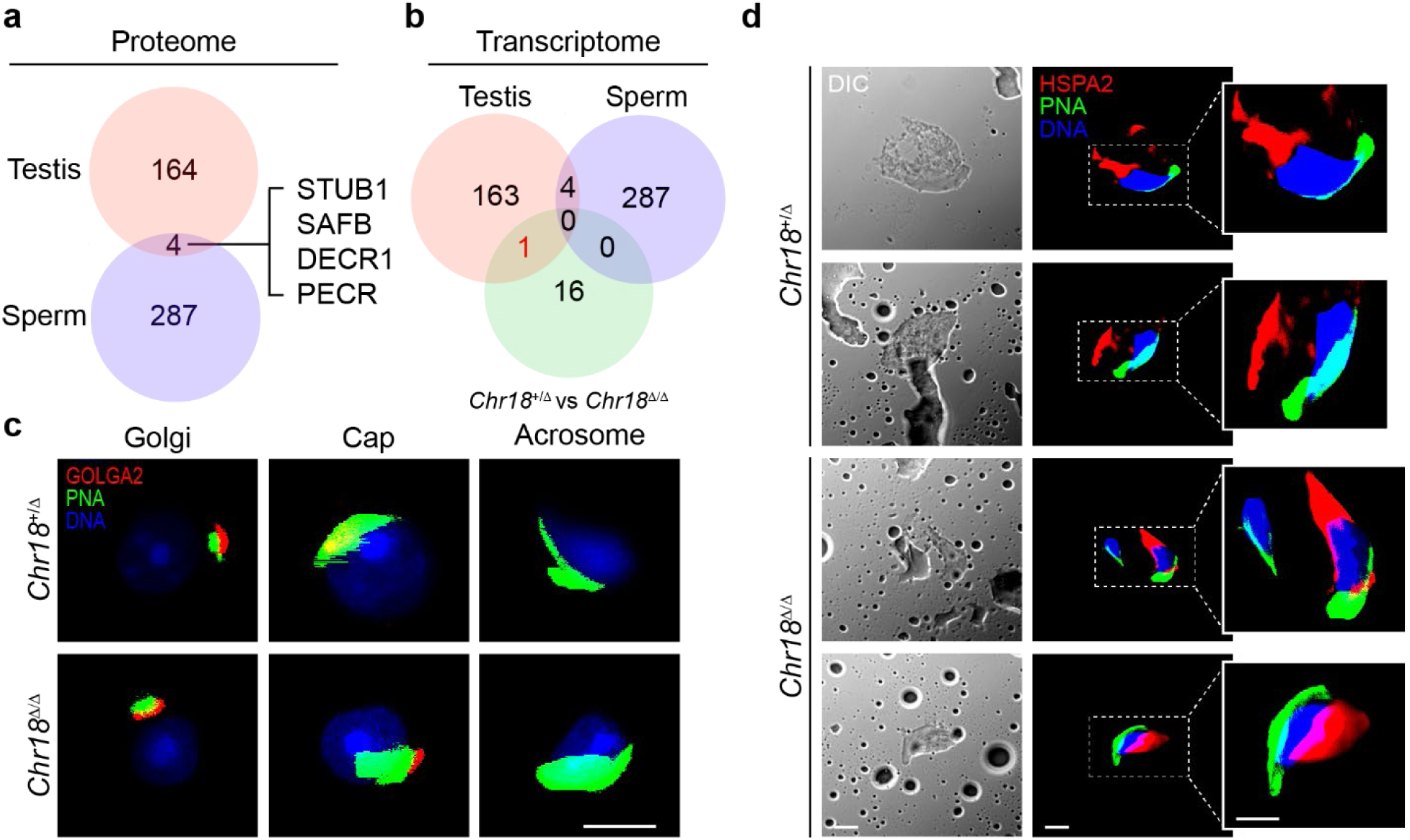
Aberrant GOLGA2 and HSPA2 expression in *Chr18^Δ/Δ^* testes and sperm. **a**, **b**, Venn diagrams depicting the overlap of altered proteins between testes and sperm proteomic profiles (**a**) and overlap of transcriptome and proteome between +/Δ and Δ/Δ mice (**b**). **c**, Representative confocal microscopic images of round and elongating spermatid heads from control and mutant mice (n =3 for each genotype). Sections were stained for GOLGA2 (red), PNA (acrosome, green) and Hoechst 33342 (nuclear DNA, blue). Scale bar, 5 μm. **d**, Representative confocal microscopic images of elongating and elongated spermatids from controls and mutant mice (n = 3). Isolated cells were stained for HSPA2 (red), PNA (acrosome, green), and Hoechst 33342 (nuclear DNA, blue) on the right and differential interference contrast (DIC) images are on the left. Scale bar, 5 μm.

**Extended Data Fig. 6.**
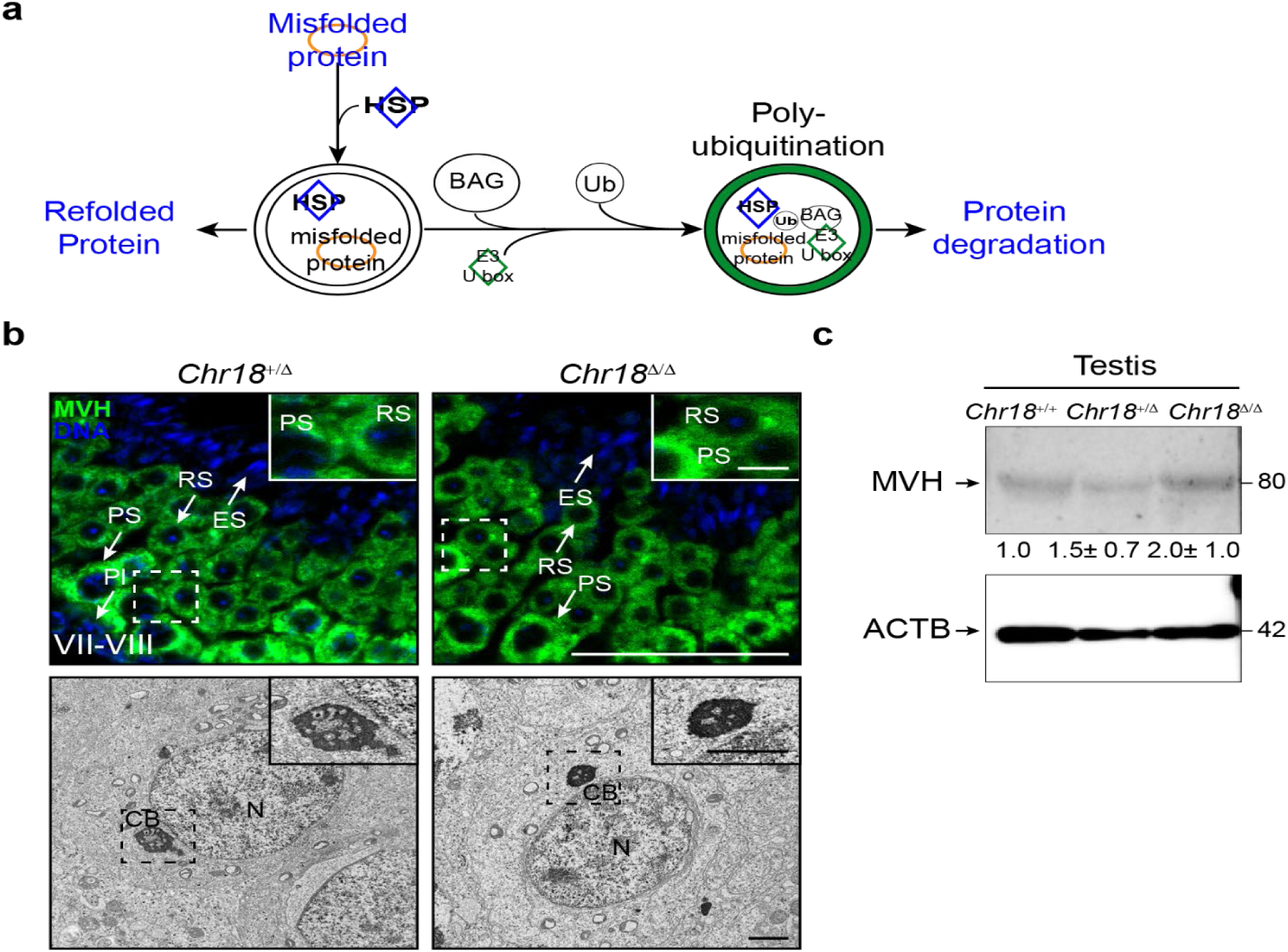
Unbalanced protein triage in *Chr18^Δ/Δ^* mice. **a**, Schematic representing protein triage. HSP functions as a chaperone to refold misfolded proteins and UPS prepares proteins for degradation. **b**, Representative confocal and TEM images of testis sections from control and mutant 8 wk/old mice (n =3 for each genotype). Sections (top) were stained for DDX4/MVH (CB, green), PNA (acrosome, green), and Hoechst 33342 (nuclear DNA, blue). Pl, preleptotene; L, leptotene; PS, pachytene spermatocyte; RS, round spermatid; ES, elongating spermatid. Scale bar, 50 μm; inset scale bar, 5 μm. The stage of seminiferous epithelium cycles was determined by morphology of spermatocytes and round spermatids stained with PNA. TEM (bottom) of round spermatids from controls and mutants (n =2). N, nucleus; CB, chromatoid body. Scale bar, 0.5 μm and the insets are 2X enlargements of the dashed box regions. **c**, Immunoblot of MVH in testes from *Chr18^+/+^*, *Chr18^+/Δ^*, and *Chr18^Δ/Δ^* (*n* = 3 for each genotype). ACTB was used as a load control. Quantification of blotting intensity for indicated proteins is shown as expression relative to *Chr 18^+/+^*controls. Data were generated from three independent experiments.

**Extended Data Fig. 7.**
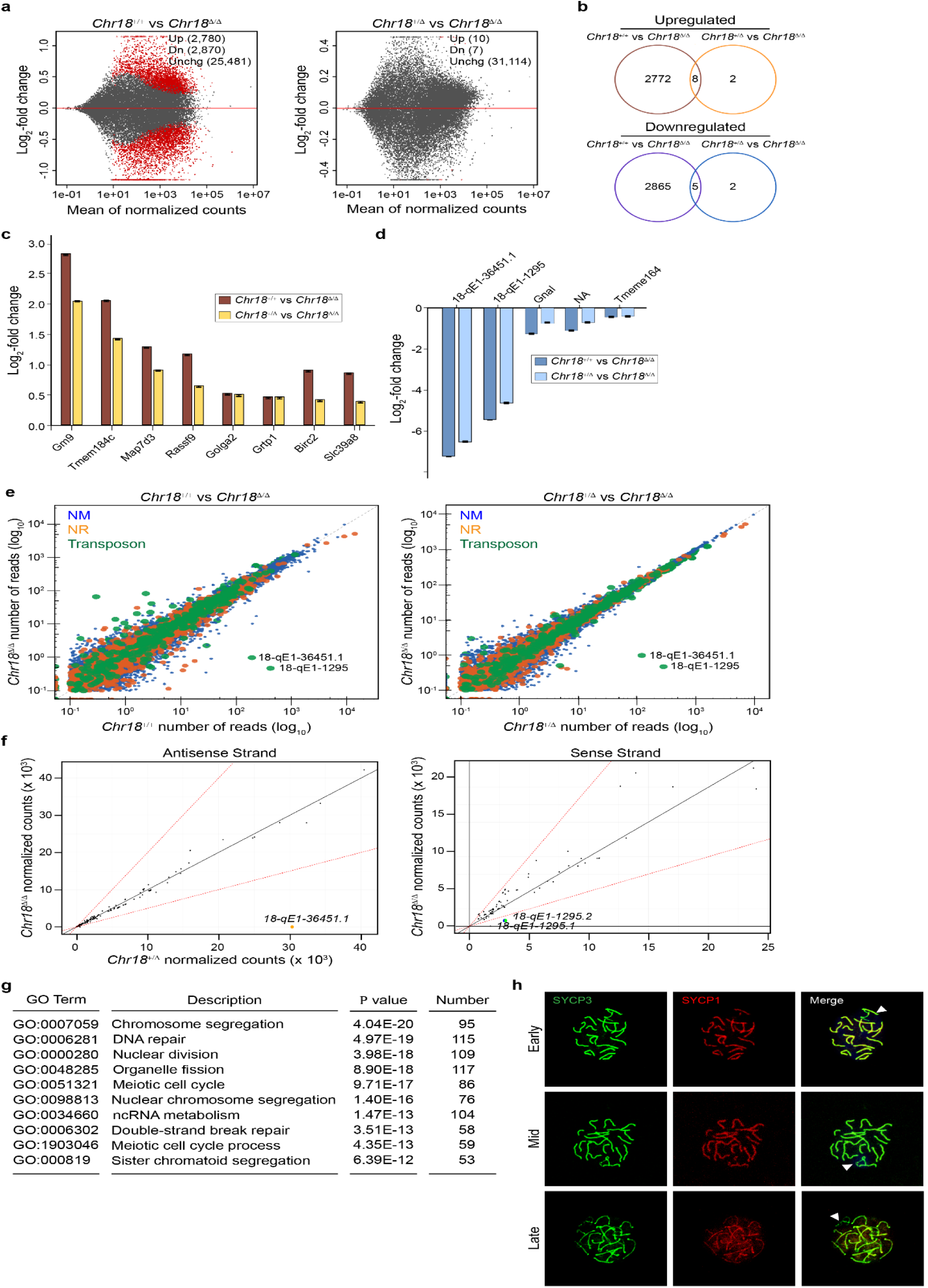
Altered abundance of transcripts in *Chr18^Δ/Δ^* mice. **a**, MA-plots of transcripts in +/+ compared to Δ/Δ (left) and +/Δ compared to Δ/Δ (right) determined by RNA-seq. The y-axis is the log_2_ fold change in expression and the x-axis is averaged expression in both genotypes. Each point represents a transcript. Transcripts with *P* adjusted values < 0.1 are colored in red. See Extended Table 3, 4. **b**, Venn diagrams depicting the overlap of up-regulated (top) and down-regulated genes (bottom) between wild-type (+/+) vs homozygous (Δ/Δ) and heterozygous (+/Δ) vs homozygous (Δ/Δ) of RNA-seq data from *Chr18^+/+^*, *Chr18^+/Δ^*, and *Chr18^Δ/Δ^* testes (*n* = 3 for each genotype) at P28 using adjusted *P* < 0.01 as the cut off. **c**, **d**, Up-regulated (**c**) and down-regulated genes (**d**) (log_2_-fold change) in Venn diagrams (**b**). See Extended Table 3, 4. **e**, Scatter plots comparing +/+ to Δ/Δ (left) and +/Δ to Δ/Δ (right) of RNA-seq reads assigned to mRNA (NM, blue); non-coding RNA (NR, orange); transposons and piRNA (green). See also Extended Table 6. **f**, Using small RNA-seq, antisense (left) and sense (right) strand piRNAs were compared between +/Δ and Δ/Δ mice. Blue and green dots, missing piRNAs on sense strand; yellow dot, missing piRNA on antisense strand. **g**, Gene ontology of differentially expressed genes after comparison of +/+ to Δ/Δ mice. See Extended Table 5. **h**, Representative confocal microscopic images of meiotic chromosome spreads of *Chr18^Δ/Δ^* pachytene (early, mid, late) spermatocytes from 4 wk/old mice. SYCP1, synaptonemal complex protein 1 (red); SYCP3, synaptonemal complex protein 3 (green). Arrowheads, sex chromosome; stage of pachytene spermatocytes was determined by desynapsis of sex chromosomes and stained with SYCP3.

**Extended Data Fig. 8.**
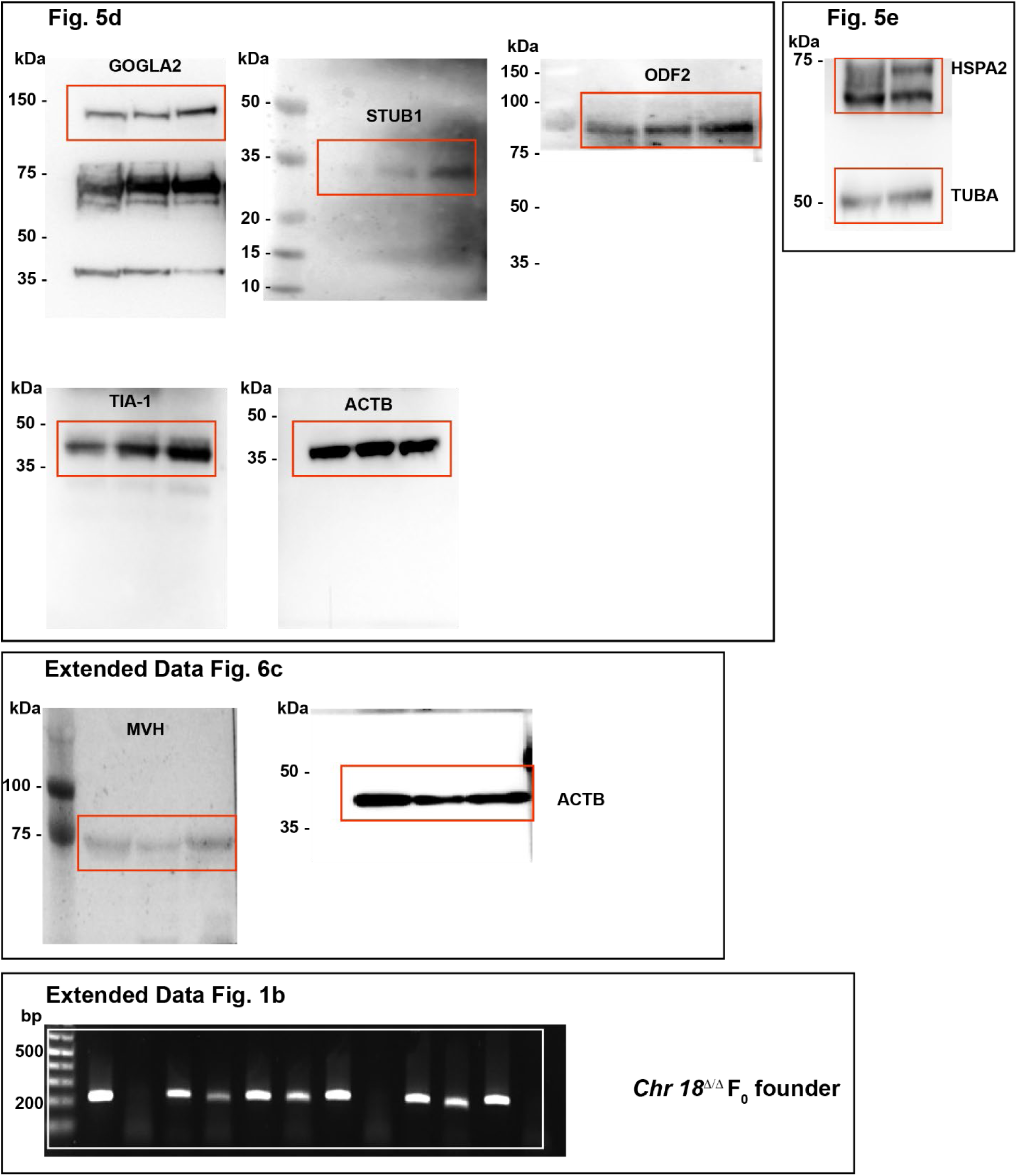
Uncropped images. Uncropped images of PCR gels and immunoblots. Rectangles indicate areas that were cropped for use in the designated figures.

## References

1. Aravin, A. et al. A novel class of small RNAs bind to MILI protein in mouse testes. Nature 442, 203–207 (2006).

2. Girard, A., Sachidanandam, R., Hannon, G.J. & Carmell, M.A. A germline-specific class of small RNAs binds mammalian Piwi proteins. Nature 442, 199–202 (2006).

3. Li, X.Z. et al. An ancient transcription factor initiates the burst of piRNA production during early meiosis in mouse testes. Mol Cell 50, 67–81 (2013).

4. Aravin, A.A. et al. A piRNA pathway primed by individual transposons is linked to de novo DNA methylation in mice. Mol Cell 31, 785–799 (2008).

5. Vagin, V.V. et al. A distinct small RNA pathway silences selfish genetic elements in the germline. Science 313, 320–324 (2006).

6. Hartig, J.V., Tomari, Y. & Forstemann, K. piRNAs--the ancient hunters of genome invaders. Genes Dev 21, 1707–1713 (2007).

7. Kuramochi-Miyagawa, S. et al. DNA methylation of retrotransposon genes is regulated by Piwi family members MILI and MIWI2 in murine fetal testes. Genes Dev 22, 908–917 (2008).

8. Houwing, S. et al. A role for Piwi and piRNAs in germ cell maintenance and transposon silencing in Zebrafish. Cell 129, 69–82 (2007).

9. Batista, P.J. et al. PRG-1 and 21U-RNAs interact to form the piRNA complex required for fertility in C. elegans. Mol Cell 31, 67–78 (2008).

10. Das, P.P. et al. Piwi and piRNAs act upstream of an endogenous siRNA pathway to suppress Tc3 transposon mobility in the Caenorhabditis elegans germline. Mol Cell 31, 79–90 (2008).

11. Gainetdinov, I., Colpan, C., Arif, A., Cecchini, K. & Zamore, P.D. A single mechanism of biogenesis, initiated and directed by PIWI proteins, explains piRNA production in most animals. Mol Cell 71, 775–790 e775 (2018).

12. Reuter, M. et al. Miwi catalysis is required for piRNA amplification-independent LINE1 transposon silencing. Nature 480, 264–267 (2011).

13. Zheng, K. & Wang, P.J. Blockade of pachytene piRNA biogenesis reveals a novel requirement for maintaining post-meiotic germline genome integrity. PLoS Genet 8, e1003038 (2012).

14. Wasik, K.A. et al. RNF17 blocks promiscuous activity of PIWI proteins in mouse testes. Genes Dev 29, 1403–1415 (2015).

15. Castaneda, J. et al. Reduced pachytene piRNAs and translation underlie spermiogenic arrest in Maelstrom mutant mice. EMBO J 33, 1999–2019 (2014).

16. Gou, L.T. et al. Pachytene piRNAs instruct massive mRNA elimination during late spermiogenesis. Cell Res 24, 680–700 (2014).

17. Goh, W.S. et al. piRNA-directed cleavage of meiotic transcripts regulates spermatogenesis. Genes Dev 29, 1032–1044 (2015).

18. Zhang, P. et al. MIWI and piRNA-mediated cleavage of messenger RNAs in mouse testes. Cell Res 25, 193–207 (2015).

19. Vourekas, A. et al. Mili and Miwi target RNA repertoire reveals piRNA biogenesis and function of Miwi in spermiogenesis. Nat Struct Mol Biol 19, 773–781 (2012).

20. Zhou, L. et al. BTBD18 regulates a subset of piRNA-generating loci through transcription elongation in mice. Dev Cell 40, 453–466 e455 (2017).

21. Yanagimachi, R. Mammalian fertilization, in The Physiology of Reproduction, Edn. 2. (eds. E. Knobil & J. Neil) 189–317 (Raven Press, New York; 1994).

22. Leblond, C.P. & Clermont, Y. Spermiogenesis of rat, mouse, hamster and guinea pig as revealed by the periodic acid-fuchsin sulfurous acid technique. Am J Anat 90, 167–215 (1952).

23. Abou-Haila, A. & Tulsiani, D.R. Mammalian sperm acrosome: formation, contents, and function. Arch Biochem Biophys 379, 173–182 (2000).

24. Kierszenbaum, A.L. & Tres, L.L. The acrosome-acroplaxome-manchette complex and the shaping of the spermatid head. Arch Histol Cytol 67, 271–284 (2004).

25. Buffone, M.G., Foster, J.A. & Gerton, G.L. The role of the acrosomal matrix in fertilization. Int J Dev Biol 52, 511–522 (2008).

26. Jin, M. et al. Most fertilizing mouse spermatozoa begin their acrosome reaction before contact with the zona pellucida during in vitro fertilization. Proc Natl Acad Sci U S A 108, 4892–4896 (2011).

27. Berruti, G. & Paiardi, C. Acrosome biogenesis: Revisiting old questions to yield new insights. Spermatogenesis 1, 95–98 (2011).

28. Ding, D. et al. TDRD5 binds piRNA precursors and selectively enhances pachytene piRNA processing in mice. Nat Commun 9, 127 (2018).

29. Bath, M.L. Inhibition of in vitro fertilizing capacity of cryopreserved mouse sperm by factors released by damaged sperm, and stimulation by glutathione. PLoS One 5, e9387 (2010).

30. Takeo, T. & Nakagata, N. Reduced glutathione enhances fertility of frozen/thawed C57BL/6 mouse sperm after exposure to methyl-beta-cyclodextrin. Biol Reprod 85, 1066–1072 (2011).

31. Ito, C. et al. Integration of the mouse sperm fertilization-related protein equatorin into the acrosome during spermatogenesis as revealed by super-resolution and immunoelectron microscopy. Cell Tissue Res 352, 739–750 (2013).

32. Roqueta-Rivera, M., Abbott, T.L., Sivaguru, M., Hess, R.A. & Nakamura, M.T. Deficiency in the omega-3 fatty acid pathway results in failure of acrosome biogenesis in mice. Biol Reprod 85, 721–732 (2011).

33. Kanemori, Y. et al. Biogenesis of sperm acrosome is regulated by pre-mRNA alternative splicing of Acrbp in the mouse. Proc Natl Acad Sci U S A 113, E3696–3705 (2016).

34. Kierszenbaum, A.L., Rivkin, E. & Tres, L.L. Acroplaxome, an F-actin-keratin-containing plate, anchors the acrosome to the nucleus during shaping of the spermatid head. Mol Biol Cell 14, 4628–4640 (2003).

35. Turner, K.J. et al. Expression cloning of a rat testicular transcript abundant in germ cells, which contains two leucine zipper motifs. Biol Reprod 57, 1223–1232 (1997).

36. Brohmann, H., Pinnecke, S. & Hoyer-Fender, S. Identification and characterization of new cDNAs encoding outer dense fiber proteins of rat sperm. J Biol Chem 272, 10327–10332 (1997).

37. Tarnasky, H. et al. Gene trap mutation of murine outer dense fiber protein-2 gene can result in sperm tail abnormalities in mice with high percentage chimaerism. BMC Dev Biol 10, 67 (2010).

38. Jiang, J. et al. CHIP is a U-box-dependent E3 ubiquitin ligase: identification of Hsc70 as a target for ubiquitylation. J Biol Chem 276, 42938–42944 (2001).

39. Kalia, S.K. et al. BAG5 inhibits parkin and enhances dopaminergic neuron degeneration. Neuron 44, 931–945 (2004).

40. Lizama, B.N. et al. Neuronal preconditioning requires the mitophagic activity of C-terminus of HSC70-Interacting protein. J Neurosci 38, 6825–6840 (2018).

41. Wang, X. et al. BAG5 protects against mitochondrial oxidative damage through regulating PINK1 degradation. PLoS One 9, e86276 (2014).

42. VanGompel, M.J. & Xu, E.Y. A novel requirement in mammalian spermatid differentiation for the DAZ-family protein Boule. Hum Mol Genet 19, 2360–2369 (2010).

43. Kim, B. & Rhee, K. BOULE, a Deleted in Azoospermia homolog, is recruited to stress granules in the mouse male germ cells. PLoS One 11, e0163015 (2016).

44. Han, F. et al. Globozoospermia and lack of acrosome formation in GM130-deficient mice. Cell Death Dis 8, e2532 (2017).

45. Dix, D.J. et al. Targeted gene disruption of Hsp70-2 results in failed meiosis, germ cell apoptosis, and male infertility. Proc Natl Acad Sci U S A 93, 3264–3268 (1996).

46. Brown, P.R., Odet, F., Bortner, C.D. & Eddy, E.M. Reporter mice express green fluorescent protein at initiation of meiosis in spermatocytes. Genesis 52, 976–984 (2014).

47. Korfanty, J. et al. Identification of a new mouse sperm acrosome-associated protein. Reproduction 143, 749–757 (2012).

48. Garrido, C. & Solary, E. A role of HSPs in apoptosis through “protein triage”? Cell Death Differ 10, 619–620 (2003).

49. Ciechanover, A. The ubiquitin-proteasome pathway: on protein death and cell life. EMBO J 17, 7151–7160 (1998).

50. Wickner, S., Maurizi, M.R. & Gottesman, S. Posttranslational quality control: folding, refolding, and degrading proteins. Science 286, 1888–1893 (1999).

51. Kim, B., Cooke, H.J. & Rhee, K. DAZL is essential for stress granule formation implicated in germ cell survival upon heat stress. Development 139, 568–578 (2012).

52. Roy, E. et al. GM130 gain-of-function induces cell pathology in a model of lysosomal storage disease. Hum Mol Genet 21, 1481–1495 (2012).

53. Lehtiniemi, T. & Kotaja, N. Germ granule-mediated RNA regulation in male germ cells. Reproduction 155, R77–R91 (2018).

54. Wu, P.H. et al. The evolutionarily conserved piRNA-producing locus pi6 is required for male mouse fertility. Nat Genet 52, 728–739 (2020).

55. Deng, W. & Lin, H. miwi, a murine homolog of piwi, encodes a cytoplasmic protein essential for spermatogenesis. Dev Cell 2, 819–830 (2002).

56. Grivna, S.T., Beyret, E., Wang, Z. & Lin, H. A novel class of small RNAs in mouse spermatogenic cells. Genes Dev 20, 1709–1714 (2006).

57. Xu, M. et al. Mice deficient for a small cluster of Piwi-interacting RNAs implicate Piwi-interacting RNAs in transposon control. Biol Reprod 79, 51–57 (2008).

58. Homolka, D. et al. PIWI slicing and RNA elements in precursors instruct directional primary piRNA biogenesis. Cell Rep 12, 418–428 (2015).

59. Dai, P. et al. A translation-activating function of MIWI/piRNA during mouse spermiogenesis. Cell 179, 1566-+ (2019).

60. Bartel, D.P. Metazoan MicroRNAs. Cell 173, 20–51 (2018).

61. Wang, H. et al. Atg7 is required for acrosome biogenesis during spermatogenesis in mice. Cell Res 24, 852–869 (2014).

62. Xiao, N. et al. PICK1 deficiency causes male infertility in mice by disrupting acrosome formation. J Clin Invest 119, 802–812 (2009).

63. Lin, Y.N., Roy, A., Yan, W., Burns, K.H. & Matzuk, M.M. Loss of zona pellucida binding proteins in the acrosomal matrix disrupts acrosome biogenesis and sperm morphogenesis. Mol Cell Biol 27, 6794–6805 (2007).

64. Funaki, T. et al. The Arf GAP SMAP2 is necessary for organized vesicle budding from the trans-Golgi network and subsequent acrosome formation in spermiogenesis. Mol Biol Cell 24, 2633–2644 (2013).

65. Tardif, S., Guyonnet, B., Cormier, N. & Cornwall, G.A. Alteration in the processing of the ACRBP/sp32 protein and sperm head/acrosome malformations in proprotein convertase 4 (PCSK4) null mice. Mol Hum Reprod 18, 298–307 (2012).

66. Yan, W., Ma, L., Burns, K.H. & Matzuk, M.M. Haploinsufficiency of kelch-like protein homolog 10 causes infertility in male mice. Proc Natl Acad Sci U S A 101, 7793–7798 (2004).

67. Pierre, V. et al. Absence of Dpy19l2, a new inner nuclear membrane protein, causes globozoospermia in mice by preventing the anchoring of the acrosome to the nucleus. Development 139, 2955–2965 (2012).

68. Kang-Decker, N., Mantchev, G.T., Juneja, S.C., McNiven, M.A. & van Deursen, J.M. Lack of acrosome formation in Hrb-deficient mice. Science 294, 1531–1533 (2001).

69. Friant, S., Meier, K.D. & Riezman, H. Increased ubiquitin-dependent degradation can replace the essential requirement for heat shock protein induction. EMBO J 22, 3783–3791 (2003).

70. Ohn, T., Kedersha, N., Hickman, T., Tisdale, S. & Anderson, P. A functional RNAi screen links O-GlcNAc modification of ribosomal proteins to stress granule and processing body assembly. Nat Cell Biol 10, 1224–1231 (2008).

71. Gerton, G.L. & Millette, C.F. Generation of flagella by cultured mouse spermatids. J Cell Biol 98, 619–628 (1984).

72. Russell, L.D., Weiss, T., Goh, J.C. & Curl, J.L. The effect of submandibular gland removal on testicular and epididymal parameters. Tissue Cell 22, 263–268 (1990).

73. Dia, F., Strange, T., Liang, J., Hamilton, J. & Berkowitz, K.M. Preparation of meiotic chromosome spreads from mouse spermatocytes. J Vis Exp (2017).

74. Roovers, E.F. et al. Piwi proteins and piRNAs in mammalian oocytes and early embryos. Cell Rep 10, 2069–2082 (2015).

75. Kim, D., Langmead, B. & Salzberg, S.L. HISAT: a fast spliced aligner with low memory requirements. Nat Methods 12, 357–360 (2015).

76. Li, H. et al. The sequence alignment/map format and SAMtools. Bioinformatics 25, 2078–2079 (2009).

77. Li, H. A statistical framework for SNP calling, mutation discovery, association mapping and population genetical parameter estimation from sequencing data. Bioinformatics 27, 2987–2993 (2011).

78. Liao, Y., Smyth, G.K. & Shi, W. The Subread aligner: fast, accurate and scalable read mapping by seed-and-vote. Nucleic Acids Res 41, e108 (2013).

79. Love, M.I., Huber, W. & Anders, S. Moderated estimation of fold change and dispersion for RNA-seq data with DESeq2. Genome Biol 15, 550 (2014).

80. Patro, R., Duggal, G., Love, M.I., Irizarry, R.A. & Kingsford, C. Salmon provides fast and bias-aware quantification of transcript expression. Nat Methods 14, 417–419 (2017).

81. Han, B.W., Wang, W., Zamore, P.D. & Weng, Z.P. piPipes: a set of pipelines for piRNA and transposon analysis via small RNA-seq, RNA-seq, degradome- and CAGE-seq, ChIP-seq and genomic DNA sequencing. Bioinformatics 31, 593–595 (2015).

82. Bao, W., Kojima, K.K. & Kohany, O. Repbase update, a database of repetitive elements in eukaryotic genomes. Mob DNA 6, 11 (2015).

83. Langmead, B. & Salzberg, S.L. Fast gapped-read alignment with Bowtie 2. Nat Methods 9, 357–359 (2012).

84. Livak, K.J. & Schmittgen, T.D. Analysis of relative gene expression data using real-time quantitative PCR and the 2(-Delta Delta C(T)) method. Methods 25, 402–408 (2001).

